# H3K27M-driven hypertranscription leads to a new targetable dependency in diffuse midline gliomas

**DOI:** 10.64898/2026.05.12.724628

**Authors:** Sharmistha Pal, Heng Wang, Jacob Geisberg, Tasmia Mirza, Khushi Kohli, Ashot Nazaretian, Truman Knowles, Christopher J. Graser, Kaitlyn Bootz, Olga Wójcikowska, Magdalena Bartyńska, Simon J. Boulton, Jayesh B. Majithiya, Helen Robinson, Graeme Smith, Charles Stiles, Dipanjan Chowdhury, Nathalie Y.R. Agar, Sabine Mueller, Franziska Michor, Daphne Haas-Kogan

## Abstract

Diffuse midline gliomas (DMGs) are driven by the H3K27M oncohistone—a challenging therapeutic target. However, conventional therapeutic modalities are never curative. Against this backdrop, we address an important unresolved question--are there H3K27M-induced oncogenic vulnerabilities that can be exploited for therapeutic benefit. We show that H3K27M induces hypertranscription, thus identifying hypertranscription as a new molecular feature of H3K27M-driven DMGs. We demonstrate this finding in genetic mouse models, human DMG cells, and primary tumor specimens. We further demonstrate that H3K27M-induced hypertranscription perturbs replication, heightens basal replication stress, and enhances sensitivity to ATR inhibition. In exploring therapeutic implications of these findings, we document brain penetrance, target engagement, and therapeutic efficacy of a clinical-stage ATR inhibitor (alnodesertib) *in vitro* and in intracranial DMG xenografts. We further demonstrate synergistic activity of alnodesertib with radiotherapy—the current standard of care for DMGs. These findings provide the mechanistic underpinning and preclinical rationale for including alnodesertib as monotherapy and in combination with radiation in clinical trials for children with H3K27M DMGs. The broad implications of our studies highlight ATR inhibition as a therapy for aggressive human cancers displaying hypertranscription.

## INTRODUCTION

Diffuse midline gliomas (DMGs) are aggressive, infiltrative brain tumors that arise in the midline structures or spinal cord in children (*1, 2*). DMGs are surgically challenging and refractory to chemotherapy. Radiotherapy is the standard of care and is the only treatment known to prolong patient survival (*3*). Only a single drug--dordaviprone (ONC201)--has received FDA approval for recurrent DMGs and no agents have been approved for newly diagnosed DMGs. Regardless of therapy, prognosis for children with DMGs is bleak. New therapeutic options are urgently needed and agents that display single-agent efficacy and also synergize with radiotherapy are particularly appealing. The prevalent oncogenic drivers for DMG are a set of recurrent mutations in histone H3.3 (*H3F3A*) or H3.1 (*HIST1H3B* and *HIST1H3C*) (*4–6*). H3K27M is the most common (80%) mutation in DMG and drives the unique DMG biology and pathology, leading to the designation “H3K27M mutant diffuse midline glioma” by the World Health Organization (*7–9*). Unlike the EGFR or PI3K oncoproteins that drive a significant cohort of adult high-grade gliomas, the H3K27M oncoproteins have no mitogenic function per se; rather, the H3K27M mutation acts as an oncogenic driver by inhibiting the EZH2 subunit of the Polycomb Repressive Complex 2 (PRC2), causing global reduction of H3K27 trimethylation (H3K27me_3_) and widespread epigenetic reprogramming. This reprogramming of the epigenetic landscape in DMGs leads to enhanced self-renewal capacity and loss of differentiation within a subpopulation of stem-like cells that drive unregulated growth of DMGs (*10*). The global loss of the transcriptional repressive H3K27me_3_ led us to hypothesize that DMGs would exhibit hypertranscription--the genome-wide increase in RNA output, discovered as a developmental phenomenon over 70 years ago and recently appreciated as a key feature of aggressive cancers (*11, 12*). Establishing whether H3K27M leads to hypertranscription in DMGs is important because such increased global transcription may lead to activation of replication stress responses and open the door to hitherto unrecognized therapeutic approaches for this deadly disease.

In this study we show that the H3K27M oncoprotein increases global transcription and perturbs replication, leading to high basal replication stress and dependency on ATR. Further, we show that patient-derived H3K27M mutated DMG cells and DMG xenografts are selectively sensitive to a clinical-stage, brain-penetrant inhibitor of ATR, with comparable *in vivo* efficacy to that observed for dordaviprone, the only FDA-approved drug for DMGs. Importantly, ATR inhibition synergizes *in vitro* and *in vivo* with radiotherapy to induce DMG cell death and prolong survival of DMG-bearing mice, paving the path for upfront clinical testing of ATR inhibition in combination with radiation in pediatric DMG trial.

## RESULTS

### H3K27M expression leads to hypertranscription

Global epigenetic reprogramming, reflected in DMGs by loss of the transcriptional repressive mark of H3K27me_3_ and increase in the transcriptional activation modification H3K27 acetylation, led us to test the hypothesis that H3K27M drives hypertranscription in DMGs. To test this hypothesis, global transcription was quantified by monitoring uptake of the uridine analog EU (5-Ethynyluridine) in isogenic human pediatric glioma cells (SF188) and isogenic murine DMG models differing only in H3K27 mutational status (H3K27M and its wild-type counterpart; fig. S1A, S1B; see methods for details). Our results show higher EU incorporation in H3K27M cells compared to their isogenic wild-type counterparts, indicating H3K27M-dependent hypertranscription (Fig. 1A, 1B). To confirm that EU uptake reflected transcription, we included a CDK9 inhibitor (inhibits transcriptional elongation) as a control, and as expected, CDK9i significantly reduced EU incorporation. We confirmed H3K27M-driven increased transcription by extending our experiments to three patient-derived H3K27M-mutated and three H3 wild-type DMG cell lines (Fig. 1C) and again found increased EU incorporation in H3K27M DMGs, diminished by CDK9i, confirming hypertranscription.

**Fig. 1.**
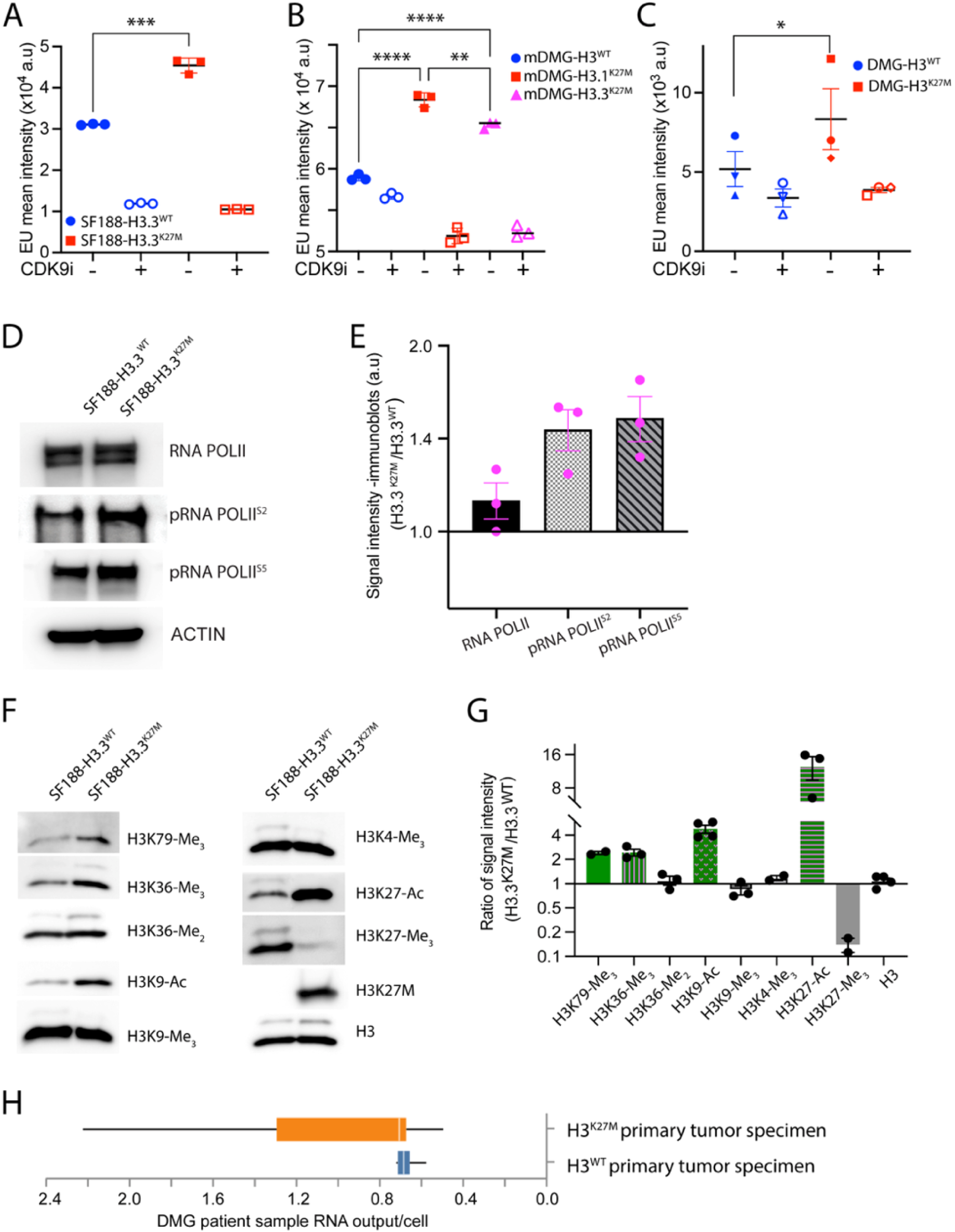
Oncohistone H3K27M induces hypertranscription. **(A-C)** Transcriptional activity in (A) isogenic SF188-H3.3^WT^ and SF188-H3.3^K27M^ cells, (B) isogenic murine DMG cells, and (C) patient-derived H3 wild-type (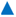 DIPG1, 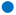 VUMC-DIPG10, 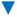 SU-DIPG48) and H3K27M-mutated (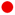 SU-DIPG4, 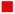 SU-DIPG13, 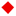 BT869) DMG cells was quantified by uridine analog (5-Ethynyluridine, EU) incorporation in flow cytometry assays. CDK9 inhibitor (atuveciclib) treatment demonstrates specificity to RNA POL2-dependent EU incorporation. **(D and E)** Western blot analysis of whole cell extracts of SF188-H3.3^WT^ and SF188-H3.3^K27M^ isogenic cells to assess RNA POL2 levels and its phosphorylation at serine 2 (S^2^-transcriptional elongation) and 5 (S^5^-initiation and promoter clearance phase). (D) A representative blot and (E) quantifications of total and phosphorylated RNA POL2 in H3K27M relative to wild-type cells (n=3). **(F and G**) Western blot analyses were performed on histone extracts from SF188-H3.3^WT^ and SF188-H3.3^K27M^ isogenic cells to assess levels of transcription-associated histone modifications using indicated antibodies. (F) Representative blots and (G) quantification (n=2-4) in H3K27M relative to wild-type cells. **(H)** Box plot of RNA output (transcriptional volume) of all single cells sequenced from DMG H3K27M mutated or wild-type primary tumor specimens.

As an independent metric of H3K27M-driven enhanced transcription we conducted two additional sets of experiments in isogenic pairs differing only in H3K27M status: (1) we assessed the status of RNA POL2 phosphorylation at serine 2 and 5 and documented increased phosphorylation of RNA POL2 at both sites—phosphorylation of which is found on actively transcribing RNA POL2 (*13*) (Fig. 1D, 1E); (2) we measured levels of overall H3K36 and H3K79 trimethylation (H3K36me_3_, H3K79me_3_), and H3K9 acetylation (H3K9-Ac), all associated with actively transcribing RNA POL2 (*14, 15*). These studies (Fig, 1F, 1G, fig. S1C) showed marked increases in levels of H3K36me_3_, H3K79me_3_, and H3K9-Ac in H3K27M cells relative to wild-type isogenic pairs. We included additional controls to document specificity of H3K27M effects on transcription-associated histone modifications. These controls included methylation levels of H3K4me_3_ (active promoters), H3K9me_3_ (heterochromatin), and H3K36me_2_ (intergenic regions). None of these methylation sites showed significant differences between isogenic pairs (Fig. 1F, 1G, fig. S1C). Finally, we documented H3K27M expression and resultant expected epigenetic changes. All H3K27M isogenic lines demonstrated H3K27M expression, loss of H3K27me_3_, and increased H3K27 acetylation (*16–18*).

Having documented that H3K27M drives hypertranscription in cell line models of DMGs—human and mouse, isogenic and non-isogenic—we asked whether the hypertranscription phenotype would be evident in primary DMG patient tumor specimens. To this end, we leveraged existing single-cell mRNA sequencing data from H3 wild-type and H3K27M mutated DMG primary tumor samples and computed transcriptional volumes at the individual cell level (*19*). Our analyses show that H3K27M mutated tumor cells had statistically significant higher levels of mRNA compared to wild-type tumor cells (p=0.011 using a linear mixed-effects model; Fig. 1H, fig. S1D, Table S1). Of note, this hypertranscription phenotype was evident in the malignant oligodendroglial precursor-like cell (OPC) population that drives DMG tumor growth and was agnostic to cell cycle phase (fig. S1E, S1F, Table S1). Thus, our data demonstrate that H3K27M enhances global transcription, leading to a hypertranscription phenotype in DMG tumors.

### H3K27M-driven hypertranscription is accompanied by increased R-loops and transcription-replication conflicts

H3K27M-driven increased transcription predicts that DMGs will exhibit higher levels of transcription-related products, specifically R-loops and transcription-replication conflicts (TRC) (*20–23*). To test this prediction, we quantified R-loops in genomic DNA from our isogenic paired cells and patient-derived DMG cell lines by immunoblotting with S9.6 antibody. The data (Fig. 2A-2D) document an increase in R-loops in H3K27M mutated cells. To document specificity of the S9.6 antibody for R-loops, we treated genomic DNA with RNASEH1 and found loss of anti-S9.6 signal, reflecting loss of R-loop RNA-DNA hybrids. We also used an alternative approach to quantitate R-loop formation by measuring binding of the V5-tagged RNASEH1^D210N^ mutant in isogenic pairs. We found increased chromatin-bound RNASEH1^D210N^ in H3K27M cells compared to isogenic wild-type counterparts, indicating significantly higher R-loops in mutant cells (fig. S2).

**Fig. 2.**
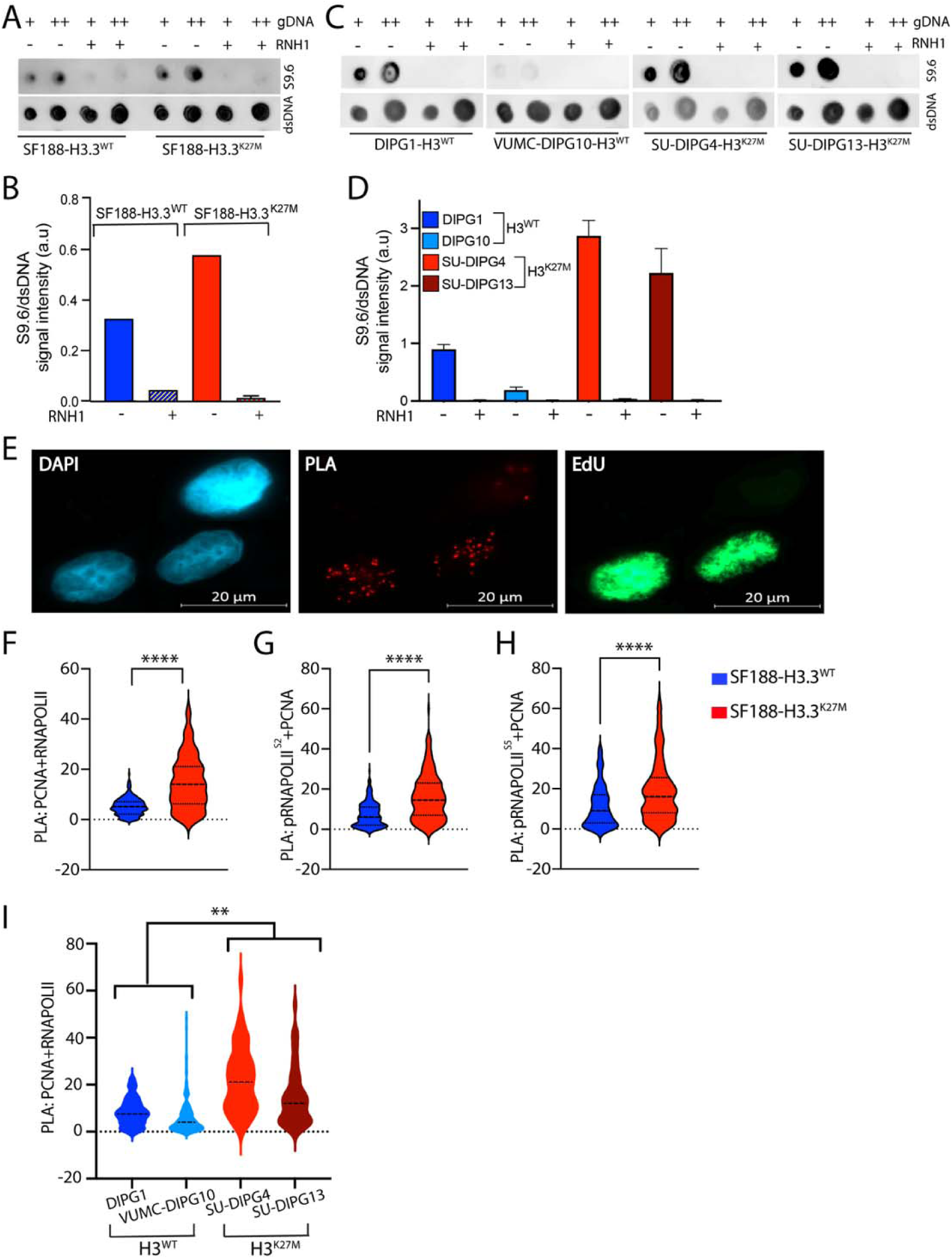
H3K27M drives increased R-loops and transcription-replication conflicts (TRC). **(A-D)** R-loops detected by anti-S9.6 dot blots of genomic DNA from (A-B) isogenic SF188-H3.3^WT^ and SF188-H3.3^K27M^ cells, and (C-D) patient-derived DMG cells. A and C show dot blots while B and D show quantification of S9.6/dsDNA signals (mean ± SEM). RNASEH1 treatment demonstrates specificity of R-loop detection; dsDNA is used as loading control. **(E-I)** Proximity ligation assays (PLA) were used to score transcription-replication conflict events in (E-H) isogenic SF188-H3.3^WT^ and SF188-H3.3^K27M^ cells, and (I) patient-derived DMG cells. Anti-PCNA (replication component) was detected in proximity to transcription components--(F and I) total RNA POL2, (G) phospho-RNA POL2^S2^, or (H) phospho-RNA POL2^S5^. EdU labeling identified S phase cells. (E) Representative images are shown. (F-I) PLA dots were quantified using Cell Profiler software, and number of dots in at least 100 EdU-positive cells is represented in violin plots.

Examining a second consequence of hypertranscription, we quantified TRC events by proximity ligation assays (PLA) in which we evaluate the proximity of PCNA (replication component) and RNA POL2 (transcription component). We observed PLA signal as distinct microscopic dots in S-phase cells and found significantly higher numbers of PLA dots (representing TRCs) in H3K27M cells relative to isogenic wild-type counterparts (Fig. 2E, 2F, 2I). PLA assays performed using antibodies specific to Ser2-phosphorylated RNA POL2 (promoter-distal region) (Fig. 2G) or Ser5-phosphorylated RNA POL2 (promoter-proximal region) (Fig. 2H) show increased TRCs in both promoter-distal and promoter-proximal regions in H3K27M cells relative to isogenic wild-type counterparts. Taken together, our results indicate that H3K27M-expressing glioma cells exhibit hypertranscription and resultant transcription-associated features (R-loops and TRCs).

### H3K27M perturbs DNA replication origin firing

A broad body of work documents that R-loops and TRCs obstruct replication fork progression (*24*), thus perturbing DNA replication and inducing replication stress. To test the prediction that H3K27M perturbs DNA replication and creates replication stress in DMG cells, we quantified overall DNA replication in the human isogenic pediatric glioma cell line pair by measuring incorporation of the deoxyuridine analog EdU. Flow cytometry assays show an increase in EdU incorporation (EdU intensity), indicating increased DNA synthesis in H3K27M cells relative to their wild-type isogenic counterparts (Fig. 3A). This is against a backdrop of similar S phase distributions between the isogenic pairs. For a critical look at cells within S phase, we divided S phase cells into --early S, mid S, and late S phase--and computed mean EdU intensities of the isogenic pairs (Fig. 3B). As expected, highest EdU incorporation occurs in mid S phase; however, regardless of S phase sub-stage (early-, mid-, or late-S phase), EdU incorporation was higher in H3K27M cells relative to wild-type cells. We extended our observations beyond isogenic cells to patient-derived DMG cell lines and documented higher EdU incorporation in H3K27M mutated compared to H3 wild-type DMGs (Fig. 3C), corroborating our findings in the isogenic cells.

**Fig. 3.**
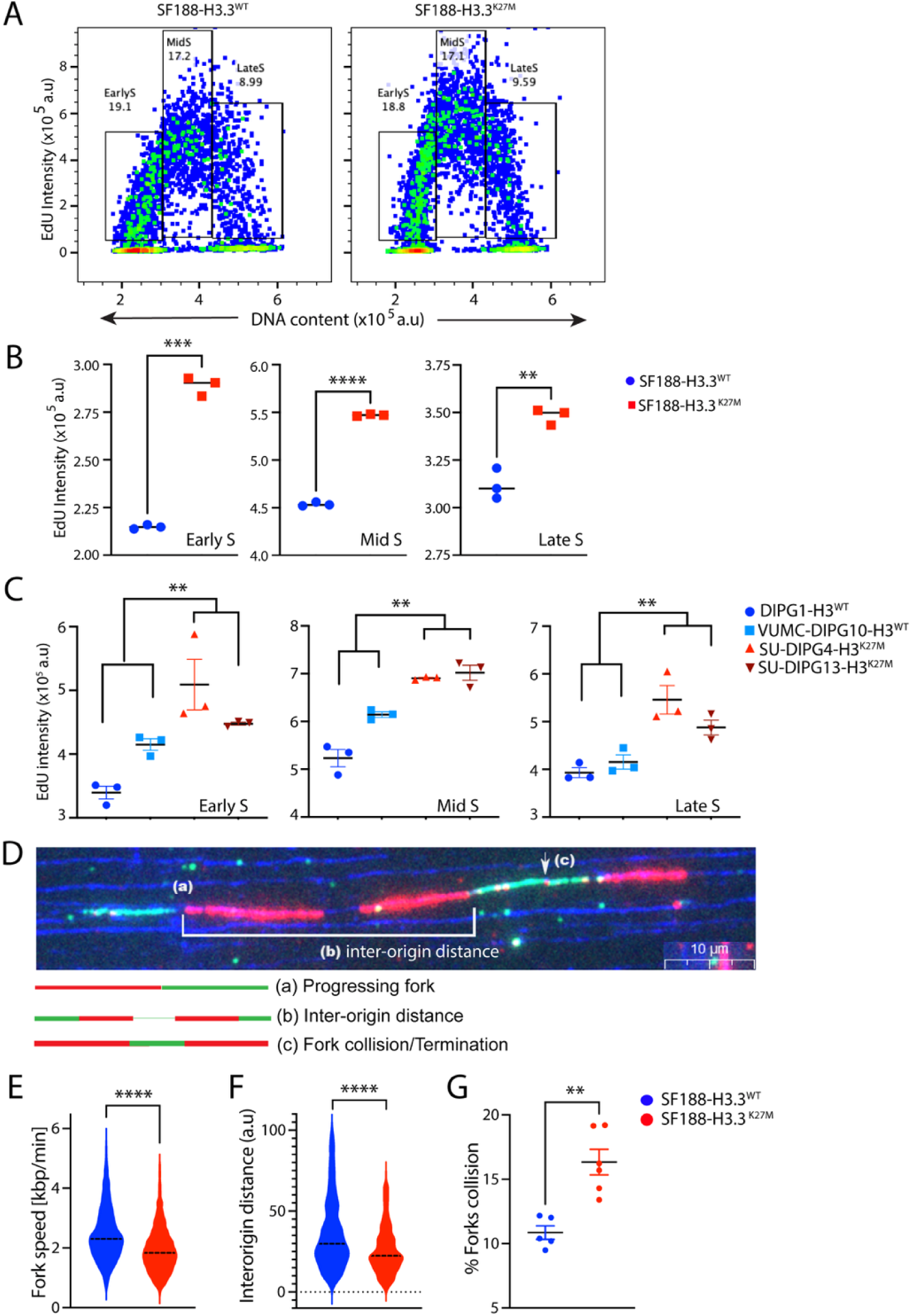
H3K27M expression disrupts DNA replication, a hallmark of replication stress. **(A-C)** DNA replication was assessed by flow cytometry in isogenic human pediatric glioma SF188-H3.3^WT^ and SF188-H3.3^K27M^ cells and patient-derived DMG cells to measure nucleotide analog (EdU) incorporation. Mean EdU incorporation was quantified in early-, mid-, and late-S phase cells. Representative flow cytometry images are shown in (A) and quantifications of EdU intensity (n=3) are plotted in (B) for the isogenic pair and in (C) for the patient-derived DMG lines (DIPG1 and VUMC-DIPG10 are H3^WT^; SU-DIPG13 and SU-DIPG4 are H3^K27M^). **(D-G)** DNA fiber assays were used to quantify replication parameters. (D) Representative image of DNA fiber assay and schematic representation of scored events. (E-G) DNA fiber assay results for isogenic SF188-H3.3^WT^ and SF188-H3.3^K27M^ cells show: (E) replication fork speed (n=3), (F) inter-origin distance (n=3), and (G) replication fork collisions (n=5-6). Data represent pooled quantification of multiple independent experiments, as indicated in parentheses.

H3K27M-dependent increased EdU incorporation documented above may be due to higher replication fork speed or greater number of replication forks, or both. To differentiate between these possibilities, we assessed replication fork dynamics at single-molecule resolution using DNA fiber assays (*25*). A representative image and schematic (Fig. 3D) show how this assay quantifies replication fork parameters, including speed, inter-origin distance, and fork collisions. As indicated, H3K27M isogenic cells exhibit slower fork speed (Fig. 3E), shorter inter-origin distances (Fig. 3F), and more fork collisions (Fig. 3G) relative to H3 wild-type cells—all features that indicate increased origin firing (*26*). Of note, H3K27M mutation did not alter replication fork stability or restart capability (fig. S3). Taken together, our results indicate that H3K27M increases DNA replication through enhanced origin firing, a phenomenon known to induce replication stress.

### H3K27M cells display higher basal replication stress and ATR dependency

H3K27M-dependent increased replication fork firing predicts that H3-mutant cells will exhibit higher basal replication stress and ATR activation. To test this hypothesis, we first measured chromatin-bound RPA in S-phase cells, reflecting single-stranded DNA levels. Whether in isogenic glioma cells (Fig. 4A, 4B, left panels) or patient-derived DMGs (Fig. 4A, 4B, right panels), higher levels of RPA-bound chromatin were documented in S-phase H3K27M cells relative to their wild-type counterparts. Since RPA-bound single-stranded DNA is a prerequisite for ATR activation, we further hypothesized that H3K27M cells would display increased basal ATR activation. Elevated RPA-bound chromatin in H3K27M cells correlated with enhanced phosphorylation of known ATR target proteins, including RPA32^S33^, CHK1^S317^, and autophosphorylation of ATR^T1989^ (Fig. 4C and fig. S4A, S4B), suggesting greater H3K27M-dependent basal ATR activation. To confirm specific activation of ATR, we examined phosphorylation of KAP1^S824^, an ATM target, and found no difference in KAP1 phosphorylation between isogenic cells (Fig. 4C). Going further, we utilized patient-derived DMG cell lines and found increased CHK1 phosphorylation in H3K27M DMGs compared to H3 wild-type DMGs (Fig. 4D). Our results indicate that cells expressing H3K27M have higher basal chromatin-bound RPA and ATR activation, both hallmarks of replication stress.

**Fig. 4.**
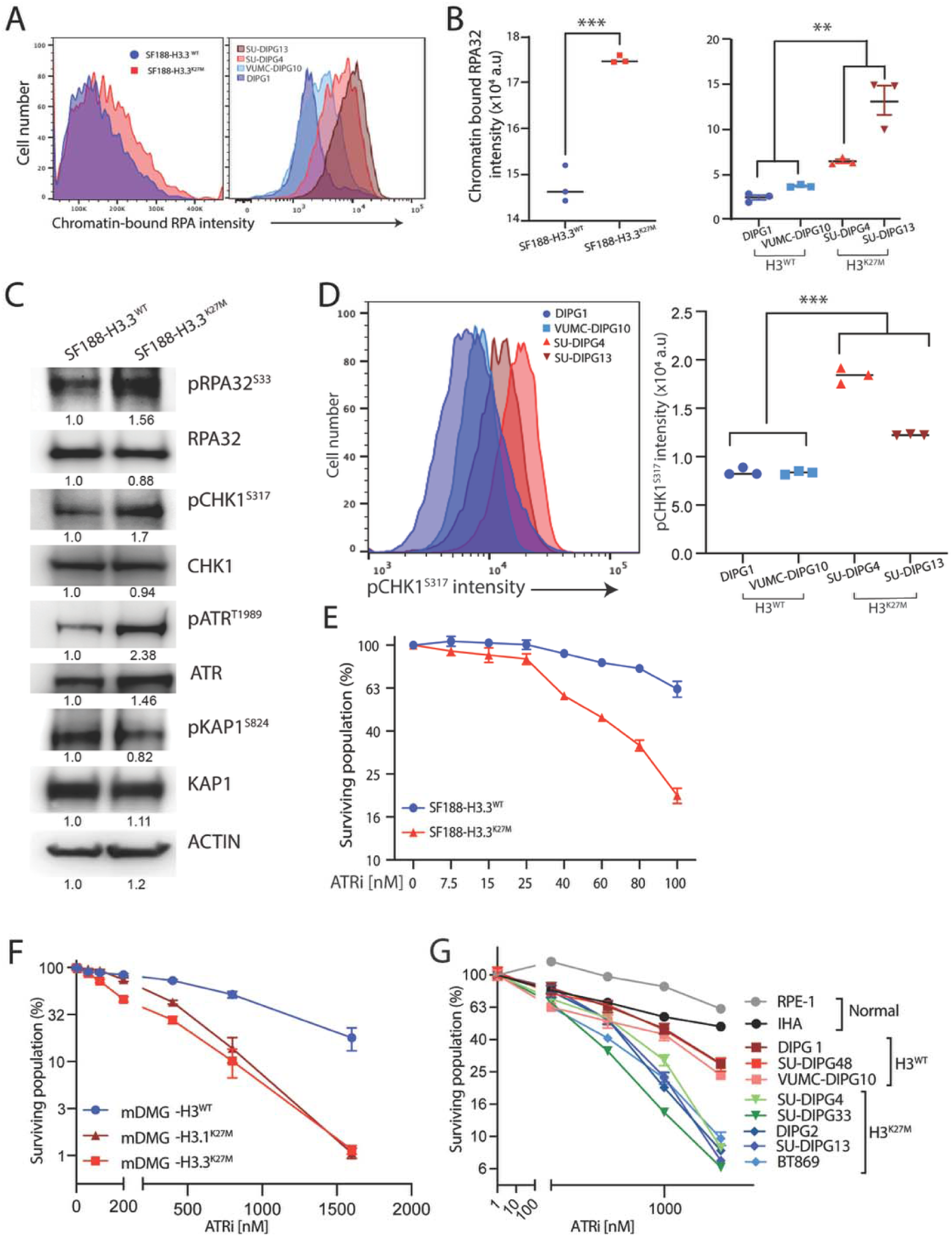
H3K27M drives higher basal replication stress and sensitivity to ATR inhibition. **(A and B)** Chromatin-bound RPA32 was measured by flow cytometry in isogenic SF188-H3.3^WT^ and SF188-H3.3^K27M^ cells and patient-derived DMG cell lines. (A) Representative histograms show chromatin-bound RPA32 in S phase cells and (B) quantification (n=3) is shown. S-phase cells were identified using EdU click-it-chemistry. **(C)** Western blot analysis was performed on isogenic SF188-H3.3^WT^ and SF188-H3.3^K27M^ whole cell extracts using indicated antibodies. **(D)** Phosphorylation of CHK1 at S317 was measured in DMG cell lines by flow cytometry. **(E)** Colony forming assays for isogenic SF188-H3.3^WT^ and SF188-H3.3^K27M^ cells were performed with varying doses of ATR inhibitor (alnodesertib); colonies were stained and quantified on day 9 (n=4, mean ± SEM). **(F and G)** Alnodesertib sensitivity was determined by Cell-titer Glo on cells from (F) isogenic murine DMG models, and (G) normal immortalized cells, along with wild-type or H3K27M-mutated patient-derived DMG cells. Graph represents surviving fractions after 6 days of alnodesertib treatment (n=6; mean ± SEM).

The findings above predict that H3K27M cells will be hyper-dependent on ATR and therefore, display increased sensitivity to ATR inhibition. Towards this goal, we tested the sensitivity of human isogenic cells differing in H3 status to the ATR inhibitor alnodesertib (*27, 28*), by colony formation assays (Fig. 4E). H3K27M cells demonstrated marked sensitivity to alnodesertib relative to their wild-type isogenic counterparts (Fig. 4E). In contrast, no differential sensitivities to ATM or DNA-PK inhibitor--ATR kinase family members (*29*)—were seen (fig. S4C, S4D). To further validate these findings, we extended our analyses to include murine isogenic DMG cell lines and a panel of patient-derived DMG cells and again found that all H3K27M DMGs exhibited enhanced sensitivity to ATR inhibition (ATRi) (Fig. 4F, 4G, and fig. S4E, S4F). Of note, sensitivity of normal immortalized human cells--astrocytes (IHA) and retinal epithelial cells (RPE-1)--to alnodesertib was limited. In sum, we find that H3K27M drives higher replication stress, enhanced reliance on ATR, and resultant sensitivity to ATR inhibition, presenting a new therapeutic intervention with an expected favorable therapeutic window.

### Mitigating hypertranscription rescues heightened replication stress and ATRi sensitivity

The relationship between H3K27M, hypertranscription, replication stress, and ATR dependence documented in Fig. 1-4 above is pervasive but correlative. To document a causal relationship between hypertranscription and heightened replication stress in H3K27M cells, we decreased transcription in H3K27M cells to levels comparable to wild-type counterparts and measured resultant levels of DNA synthesis, replication stress, and ATRi sensitivity. To decrease overall transcription, we targeted CDK8--a non-essential gene in the mediator complex—which, of note, only minimally affected cell viability (*30*) (fig. S5A). CDK8 inhibition (CDK8i) lowered transcription in H3K27M cells to levels observed in their H3 wild-type isogenic counterparts (Fig. 5A). Next, we examined whether mitigating H3K27M-driven hypertranscription with CDK8i would lower DNA synthesis levels to those observed in isogenic wild-type cells. The increased DNA synthesis we had observed in H3K27M cells (as measured by EdU incorporation) was abolished after CDK8i in all sub-stages of S phases (early-, mid-, late-S phase) (Fig. 5B). Of note, CDK8i did not significantly alter cell cycle distribution of either H3K27M or wild-type cells (fig. S5B).

**Fig. 5.**
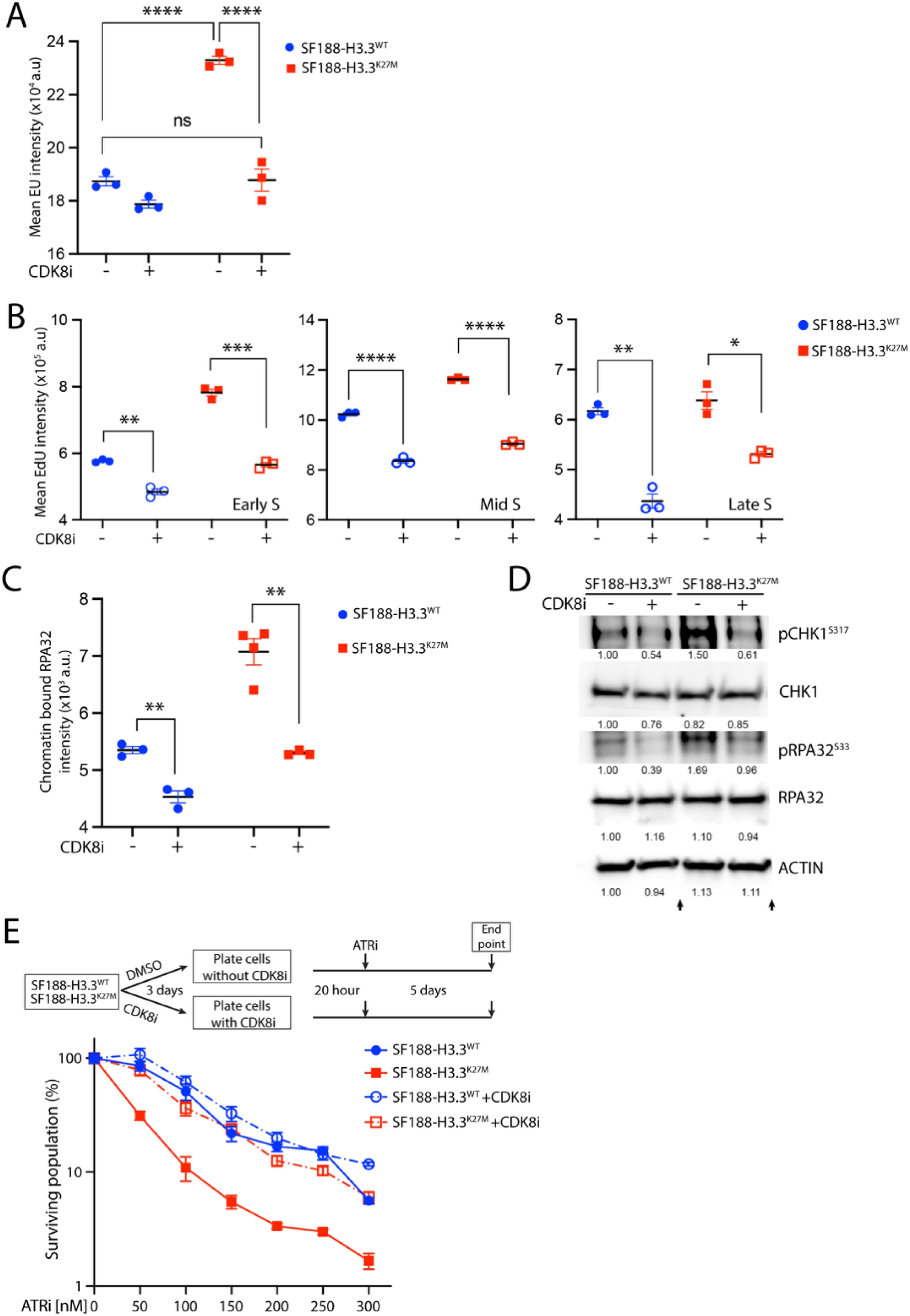
Reduced transcription rescues H3K27M-driven replication stress and hypersensitivity to ATR inhibition. **(A)** Isogenic SF188-H3.3^WT^ and SF188-H3.3^K27M^ cells were treated with MSC2530818 (CDK8 inhibitor) and transcriptional activity was measured by uridine analog (5-Ethynyluridine, EU) incorporation. **(B)** DNA replication was assessed in isogenic SF188-H3.3^WT^ and SF188-H3.3^K27M^ cells by measuring nucleotide analog (EdU) incorporation with or without CDK8i. Mean EdU incorporation was quantified in early-, mid-, and late-S phase cells. **(C and D)** Replication stress was assessed by measuring (C) chromatin-bound RPA by flow cytometry and (D) ATR-mediated phosphorylation of CHK1 and RPA by western blots in cells that were treated with either DMSO or CDK8i (arrows indicate cropping of western blots for simplicity). **(E)** SF188-H3.3^WT^ and SF188-H3.3^K27M^ cells were treated with neoadjuvant and concurrent CDK8i or DMSO (as control), as well as with increasing doses of alnodesertib (ATRi; see schematic) before cell viability was measured by CellTiter-Glo (mean ± SEM; n=4). For all the above assays, cells were treated with 1 μM CDK8i for 72 hours.

Decreased DNA synthesis following inhibition of transcription (by CDK8i) predicts that CDK8i in H3K27M cells would also reduce basal replication stress and ATR activation. We found that indeed CDK8i in H3 mutant cells lowered basal replication stress, as measured by chromatin-bound RPA (Fig. 5C), and ATR activation, measured by phosphorylation of ATR targets--RPA^S33^ and CHK1^S317^ (Fig. 5D). CDK8i also lowered DNA synthesis, replication stress, and ATR activation in H3 wild-type cells but to a significantly lesser degree.

The final prediction is that reducing transcription would rescue the hyper-sensitivity of H3K27M cells to ATRi. To test this prediction, we examined ATRi sensitivity after CDK8 inhibition and found that H3K27M isogenic cells were no longer hyper-sensitive to ATR inhibition compared to wild-type cells (Fig. 5E). Collectively, these results document a causal relationship between H3K27M-driven hypertranscription, replication stress, ATR activation and sensitivity to ATRi.

### Radiation exacerbates replication stress and synergizes with ATR inhibition

Although never curative, radiotherapy extends survival and is standard of care for children with DMGs. Since radiation also increases replication stress (*31, 32*), we asked whether ATR inhibition would synergize with radiation in killing DMG cells. As experimental endpoints, we quantified replication stress (as measured by chromatin-bound RPA) and consequences of high replication stress, including DNA damage as measured by g-H2AX, genomic instability as measured by micronuclei formation, and apoptosis as measured by caspase 3/7 activity. As indicated in Fig. 6A-6D and fig. S6, concurrent ATRi and radiation treatment showed synergistic increases in chromatin-bound RPA, g-H2AX, micronuclei positive cells, and caspase 3/7 activity. Of note, ATRi and radiotherapy altered cell cycle distribution as expected and similarly in H3K27M and H3 wild-type isogenic cells (fig. S6A). To quantify a synergistic relationship between ATRi and radiation, we used the Combenefit interactive platform (*33*) to analyze synergism-antagonism of the two therapeutic modalities in human pediatric glioma isogenic cells differing in H3 status and patient-derived DMG cells. We document synergistic cell killing by ATRi and radiotherapy in all cells tested with more pronounced synergy in H3K27M expressing cells (Fig. 6E, 6F).

**Fig. 6.**
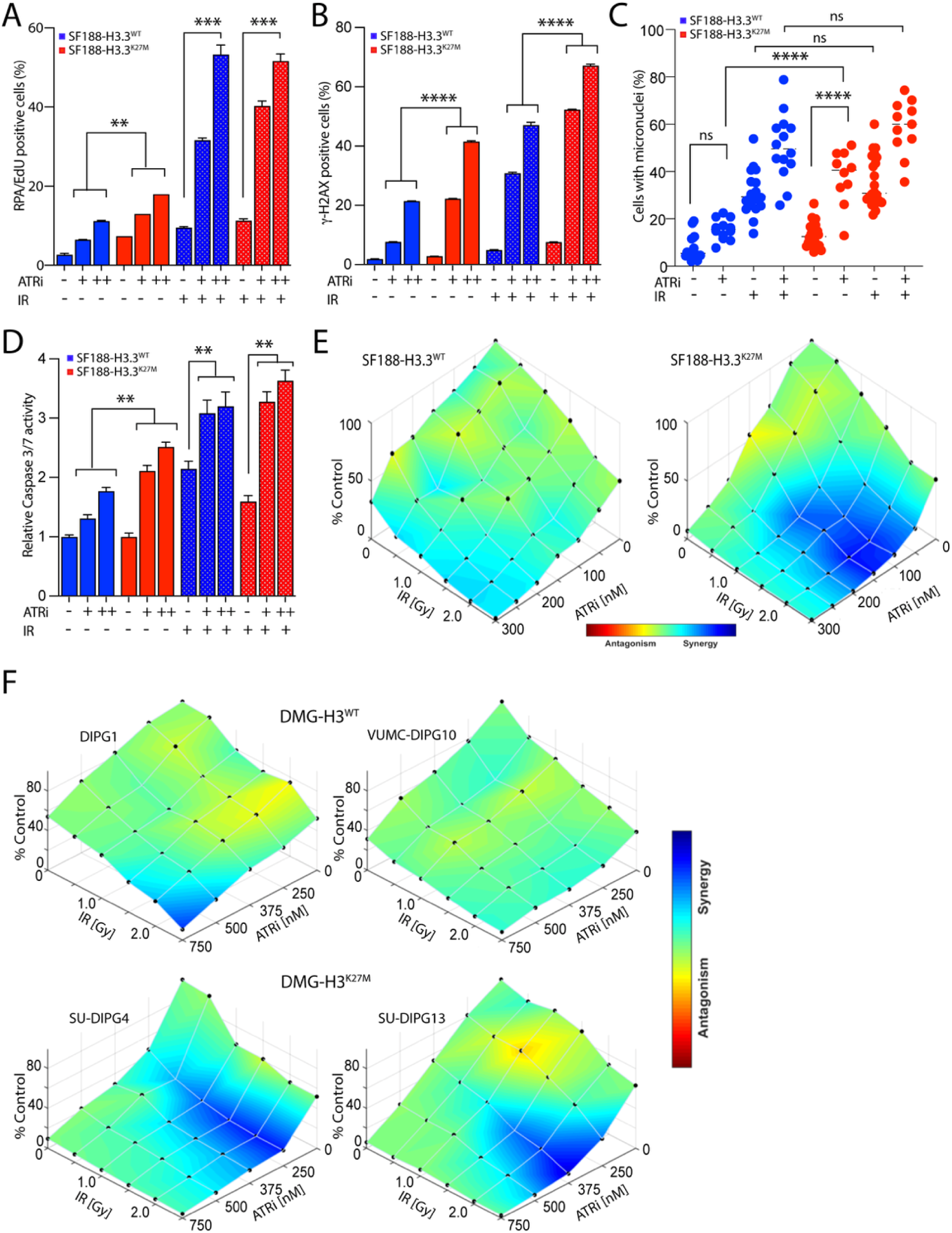
Radiation enhances replication stress and synergizes with ATR inhibition, most prominently in H3K27M cells. **(A-D)** SF188-H3.3^WT^ and SF188-H3.3^K27M^ cells were treated with ATRi and radiation as monotherapies or in combination to measure (A) chromatin-bound RPA32 (ATRi: 500 and 1000 nM; IR: 2 Gy; 24 hours; n=3); (B) DNA damage based on γ-H2AX (ATRi: 500 and 1000 nM; IR: 2 Gy; 48 hours; n=3); (C) micronuclei positive cells (ATRi: 500nM; IR: 2 Gy; 24 hours; n=3), and (D) induction of apoptosis based on CASPASE 3/7 activity (ATRi: 250 and 500 nM; IR: 1.5 Gy; 72 hours; n=3). Bar charts represent the mean ± SEM, and in panel C, each dot represents the data from a unique field of view with > 20 cells. **(E-F)** Synergy-antagonism of radiation and ATRi assessed by cell viability after 6 days of treatment (analyzed by Combenefit software) in isogenic (E) SF188-H3.3^WT^ and SF188-H3.3^K27M^ cells and (F) patient-derived DMG cell lines (n=3).

### ATR inhibitor, alnodesertib, improves survival of mice bearing DMG xenografts

The ability to cross the blood-brain barrier (BBB) is important for a drug’s therapeutic efficacy against brain tumors. Accordingly, we used matrix-assisted laser desorption/ionization mass spectroscopy imaging (MALDI-MSI) to visualize (Fig. 7A) and quantify (Fig. 7B, fig. S7A, S7B) accumulation of alnodesertib in mice brains bearing DMG tumors. MALDI-MSI analysis showed distribution of alnodesertib throughout the brain of murine DMG xenografts with higher accumulation in DMG tumor regions (Fig. 7A). Quantification indicates micromolar levels of alnodesertib in tumor regions – a level that would be expected to exhibit anti-tumor activity and indeed, alnodesertib significantly prolonged the survival of mice with H3K27M mutated SU-DIPG13-P* DMG xenografts, with 3 mice surviving to study termination (Fig. 7C). Beyond anti-tumor activity, we documented intracranial target engagement of alnodesertib by analyzing expression of the pharmacodynamic markers g-H2AX and pKAP1^S824^, in tumor tissues harvested from vehicle- or alnodesertib-treated mice (Fig. 7D, 7E). Increases in DNA damage, as measured by γ-H2AX, and phosphorylation of KAP1^S824^ are both well-established pharmacodynamic biomarkers of ATR inhibition (*34, 35*). Immunostaining showed an increased percentage of γ-H2AX- and pKAP1^S824^-positive cells in tumor regions from alnodesertib-treated mice (Fig. 7D, 7E). Finally, we asked whether the combination of ATRi and radiotherapy *in vivo* recapitulates synergistic anti-DMG activity seen *in vitro*. To this end, we treated mice bearing SU-DIPG13P* DMG xenografts with placebo (vehicle), alnodesertib (dosing at half the MTD), g-radiation, or combinations thereof. These *in vivo* randomized studies showed significantly prolonged overall survival after combination therapy compared to either alnodesertib or radiation monotherapy, which each showed minimal efficacy at these low single-agent doses (Fig. 7F).

**Fig. 7.**
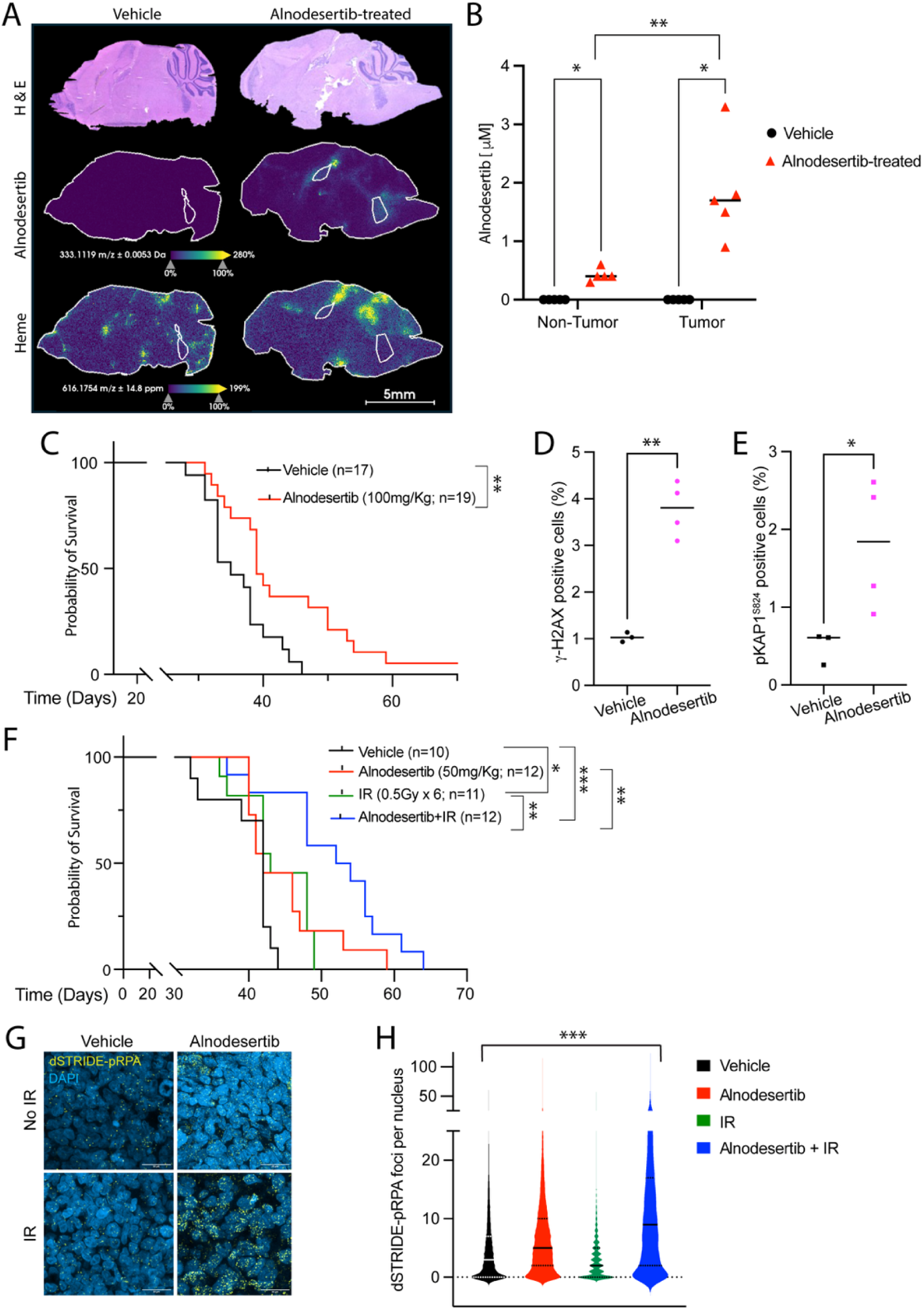
Alnodesertib (ATR inhibitor) is brain-penetrant and prolongs survival of mice bearing DMG xenografts. **(A and B)** MALDI-MSI analysis was performed on brain sections of mice bearing DIPG1 DMG orthotopic tumors after oral dosing with alnodesertib or vehicle control (100 mg/Kg, twice daily, 3 days). (A) Representative images to visualize tumor (H&E), alnodesertib, and vasculature (heme). (B) Quantification of alnodesertib (n=5) in tumor and non-tumor regions. **(C)** SU-DIPG13-P* DMG orthotopic tumor-bearing mice were randomized to alnodesertib or vehicle control and treated for 8 weeks as indicated. Overall survival (OS) analyzed by Kaplan-Meier curves. **(D and E)** Quantification of *in vivo* pharmacodynamic markers, (D) γ-H2AX and (E) pKAP1^S824^, in intracranial tumor regions of mice in randomized study (alnodesertib vs vehicle from panel C). **(F)** Overall survival analysis by Kaplan-Meier estimates of mice bearing SU-DIPG-13P* orthotopic DMG tumors and randomized to receive radiation, alnodesertib, radiation+alnodesertib combination, or vehicle control as indicated. **(G)** Representative images and (H) pooled quantification of *in situ* co-localized DSB and pRPA foci (dSTRIDE-pRPA) in tumor sections from mice (n=2-3) treated in (F), the four-arm *in vivo* experiment above.

A final set of studies showed that enhanced survival of mice in the combination arm correlated with enhanced replication stress within the tumor xenografts. To measure replication stress *in vivo*, we used the dSTRIDE-pRPA assay that quantifies replication fork collapse *in situ* by detecting phosphorylated RPA at or near a DNA double-strand break (DSB). We observed more foci of co-localized DSB and pRPA in tumor tissues from mice treated with a combination of alnodesertib and radiation compared to vehicle- or monotherapy-treated mice. Our results indicate that the survival advantage offered by alnodesertib plus radiation therapy is also associated with significantly greater *in vivo* replication stress in combination-treated K27M-mutated DMG xenografts.

## DISCUSSION

The H3K27M mutation (a change from lysine to methionine at position 27 on histone H3) defines the most aggressive pediatric brain tumor entity—“diffuse midline glioma, H3 K27-altered.” H3K27M is an oncohistone that drives tumorigenesis through global epigenome and chromatin reprogramming (*36, 37*). Prior studies have focused on H3K27M-associated changes in transcriptional programs (*38–40*); however, in this study, we specifically focused on the impact of H3K27M on global transcription because we hypothesized that such alterations would help identify hitherto unknown therapeutic approaches for the treatment of DMGs.

We show here that H3K27M leads to a global increase in transcription, also termed hypertranscription, in patient-derived DMG cell lines and primary DMG tumor samples. We thus identify hypertranscription as a new molecular feature of H3K27M-driven DMGs (*14, 15*). This H3K27M-driven hypertranscription creates replication stress, resulting in hyper-dependency on ATR--a druggable protein kinase that is required during S-phase in cells experiencing replication stress. We harnessed radiation, the standard of care treatment for DMGs, to produce yet more replication stress, resulting in marked synergy between ATR inhibition and radiation, evident most dramatically in H3K27M mutant cells. Collectively, the biology of ATR, and the studies described here, show that H3K27M-induced hypertranscription creates a DMG vulnerability with clinical promise.

Hypertranscription has never been reported as an H3K27M-mutant associated phenotype, and our findings resonate with previous reports by Krug et al., of pervasive deposition of the transcription activation mark H3K27 acetylation across the genome and increased expression of repetitive elements (*17*). In fact, hypertranscription is increasingly recognized as a feature of aggressive human cancers and has garnered interest as a potential therapeutic target for multiple cancers (*11, 41, 42*). In this study, we chose to focus on ATR inhibition as a promising target for DMGs; however, the finding of H3K27M-associated hypertranscription and increased basal replication stress has broader implications, as it opens the door to additional therapeutic approaches for DMGs. One example of a rational combination treatment supported by our findings is the combined administration of an ATR inhibitor along with a DHODH inhibitor (inhibits *de novo* pyrimidine biosynthesis), as we have previously reported that inhibition of *de novo* pyrimidine biosynthesis in DMGs exacerbates replication stress and ATR dependency (*43*).

Our study results and interpretation have some limitations. First, although alnodesertib extends survival of mice bearing H3K27M-mutated DMG xenografts, the majority of mice were not cured. Achieving cure for children with DMGs is a lofty goal that will likely require combination therapies that we will explore in future studies. Rational biology-driven, as opposed to empiric, combinations are our goal, but have not always been the norm in the field of pediatric and oncology. Second, our interpretation of the results implies broad application of ATR inhibition across a multitude of aggressive cancers that display hypertranscription and resultant replication stress. For example, in a phase 2 trial, Thomas et al. identify DNA replication stress as a therapeutic vulnerability in small cell neuroendocrine cancers those tumors with enhanced replication stress were more likely to respond to ATR and topoisomerase I inhibition (*44*) However, we do not explicitly show broad applicability of hypertranscription-dependent replication stress as a biomarker for ATR inhibition response, and such wide applications await additional clinical trials.

Our results are sufficient to introduce alnodesertib into a “basket” trial of novel agents combined with radiation for patients with DMG. Specifically, we plan to incorporate alnodesertib into the PNOC022 trial, known as the Diffuse Midline Glioma Adaptive Combinatorial Trial (DMG-ACT), an international phase II/III “basket” trial (NCT05009992) evaluating combination therapies for patients with DMGs. However, additional steps will aid in the applications of our findings in the clinic. Specifically, although H3K27M drives hypertranscription and predicts sensitivity to ATR, we see a range of global transcriptional output in our primary tumor specimens. Therefore, although H3K37M would be a predictive biomarker of response to ATR inhibition, individual tumor quantification of hypertranscription and/or replication stress may provide additional predictive value for individual patients. We are developing a formalin fixed paraffin embedded (FFPE) tissue compatible *in situ* assay to co-detect H3K27M expression and quantify basal replication stress in DMG diagnostic biopsies. The best setting in which to test this assay and its predictive value will be under the auspices of our planned clinical trial.

In early phase studies in adults, alnodesertib has shown significant anti-tumor activity in ATM-deficient advanced/metastatic solid tumors with broad histologies (NCT04657068). Alnodesertib has recently been granted Fast Track Designation by the U.S. Food and Drug Administration (FDA), in combination with low-dose irinotecan for the treatment of adult patients with ATM-negative metastatic colorectal cancer (mCRC) (*45*). The encouraging clinical results described above lend promise to a clinical trial for alnodesertib and radiation in DMGs based on an interesting parallel, evident in our results. As ATM-deficient tumors display high basal replication stress (*46*), so do H3K27M DMGs (this study). And as low-dose irinotecan enhances this high basal replication stress, so does radiotherapy in DMGs. Thus, our results provide the mechanistic underpinnings and pre-clinical rationale for a study of ATR inhibition in combination with radiotherapy for the treatment of DMGs.

## MATERIALS AND METHODS

### Antibodies, drugs and chemicals

All antibodies used in this study are listed in supplementary Table S2. Alnodesertib (ATR inhibitor) was obtained from Artios Pharma Limited under a material transfer agreement. Atuveciclib (CDK9 inhibitor) and MSC2530818 (CDK8 inhibitor) were purchased from Selleck Chemicals. 5-ethynyl uridine (EU) and hydroxyurea (HU) were purchased from Sigma-Aldrich.

### Cell lines and culture conditions

SF188 was obtained from the Brain Tumor Research Center (BTRC) Tissue Bank at the University of California, San Francisco (UCSF); patient-derived DMG cell lines--BT869, SU-DIPG13 were obtained from The Center for Patient Derived Models (CPDM) at Dana-Farber Cancer Institute (DFCI); CCHMC-DIPG1 and CCHMC-DIPG2, referred to as DIPG1 and DIPG2 were obtained from Dr. Rachid Drissi at Cincinnati Children’s Hospital Medical Center; VUMC-DIPG10 were obtained from Dr. Esther Hulleman at Princess Máxima Center for Pediatric Oncology; SU-DIPG13P*, SU-DIPG4, SU-DIPG33, and SU-DIPG48 lines were provided by Dr. Michelle Monje at Stanford University. Murine isogenic DMG cell lines established from primary DMG mouse models were provided by Dr. Timothy Phoenix at Cincinnati Children’s Hospital Medical Center (*47*). Immortalized human astrocytes (IHA; SVG p12) and retinal epithelial cells (RPE-1; CRL-4000) were purchased from ATCC.

SF188 and isogenic SF188-H3.3^WT^, SF188-H3.3^K27M^ cells, RPE-1, and IHA were cultured as adherent cells in DMEM supplemented with 10% FBS and penicillin-streptomycin (1X). Patient-derived DMG cell lines were cultured in 1:1 Neurobasal media and DMEM F-12 with MEM sodium pyruvate solution (1 mM), MEM non-essential amino acids (1X), GlutaMAX-I supplement (1X), HEPES buffer (10 mM), penicillin-streptomycin (1X), B27 supplement (1X), human-EGF (20 ng/ml), human-FGF2 (20 ng/ml), PDGF-AA (10 ng/ml), PDGF-BB (10 ng/ml), and 0.0002% heparin. Murine isogenic DMG cell lines referred to as mDMG-H3^WT^, mDMG-H3.1^K27M,^ and mDMG-H3.3^K27M^ were grown in Neurobasal media supplemented with GlutaMAX supplement (1X), penicillin-streptomycin (1X), N2 supplement (1X), B27 supplement (1X), human-EGF (20 ng/ml), human-FGF2 (20 ng/ml), and 0.0002% heparin. All human and murine DMG cell lines were grown as neurospheres except SU-DIPG4 and SU-DIPG33, which were cultured as adherent cells. All cells were cultured aseptically at 37°C in humidified incubators with 5% CO2.

### Establishment of genetically engineered cell lines

Isogenic human pediatric SF188 glioma cells were generated by knocking out *H3F3A* and exogenously expressing HA-Flag tagged-H3.3 that was either wild-type or K27M mutated. 293T cells were transfected with pLentiCRISPRv2-*H3F3A*gRNA1 (CAGACTGCCCGC AAATCGAC; Genscript), psPAX2 (Addgene #12260) and the envelope (pMD2.G; Addgene #12259) using lipofeactamine 3000 as per the manufacturer’s instructions to package lentiviral particles. SF188 cells were transduced with the *H3F3A* targeting gRNA viral particles, and after 48 hours, cells were selected with 2 mg/ml puromycin for 10 days. *H3F3A* knockout (*H3F3A* KO) was confirmed by western blot analysis of H3.3 protein levels on the pooled population, and then single cell clones of *H3F3A* KO were derived and confirmed by sequencing of PCR amplified *H3F3A* gene fragment encompassing the Cas9 site. Next, to express wild-type or K27M-mutated H3.3, SF188/*H3F3A* KO cells were transduced with lentiviral particles of either pCDH-EFI-MCS-IRES-PURO-H3.3^WT^ or pCDH-EFI-MCS-IRES-PURO-H3.3^K27M^ to establish isogenic SF188-H3.3^WT^ and SF188-H3.3^K27M^ cells and grown for about 10 passages before any experiments were performed. To express V5-tagged-RNASEH1^D210N^ in SF188-H3.3^WT^ and SF188-H3.3^K27M^, cells were transfected with 5 μg of ppyCAG_RNaseH1_D210N (*48*) (Addgene Plasmid #111904) using lipofectamine 2000 as per the manufacturer’s instructions to express V5-tagged RNASEH1^D210N^. Cells expressing V5-RNASEH1^D210N^ were selected with hygromycin (4 mg/ml) for 2 weeks before expression was confirmed by western blotting.

### Cell cycle, DNA synthesis, and transcription assessments

To determine cell cycle profiles and measure rates of DNA synthesis, EdU (10 mM) was added to the media of actively growing cells (60-70% confluency) for 1 hour before EdU was labeled using click-it-chemistry. To measure global transcription, actively growing cells were incubated with EU (0.2 mM) for 1 hour before EU was labeled using click-it chemistry. To perform click-it chemistry, cells were collected by trypsinization, fixed in 4% paraformaldehyde for 15 minutes, permeabilized with 0.5% Triton X-100 in 1X PBS for 10 minutes, and resuspended in 100 ml of 1X saponin buffer (ThermoScientific, J63209-AK) for 5 minutes. 250 ml of click-it reaction solution (ApexBio, K1179) prepared as per instructions were added, and cells were incubated at room temperature for 30 minutes before they were washed twice with washing buffer (1X PBS containing 0.5 mg/ml bovine serum albumin (BSA)) and resuspended in 250 ml of washing buffer with 1:500 diluted Hoescht33342 stain (20 mM). Cells were collected by flow cytometry on CYTOFLEX S; data were analyzed using FlowJo software. All the experiments were performed in triplicate. For analysis of DNA synthesis rates, S-phase cells were split into early-, mid- and late-S phase before mean EdU intensity was quantified.

### Whole cell lysate, total histone extraction, and western blotting

To prepare whole cell lysates, actively growing 8-10x10^6^ cells were harvested by trypsinization and lysed in UTB buffer (50 mM Tris-HCl [pH 7.5], 8 M Urea, 150 mM β-mercaptoethanol) supplemented with complete mini protease inhibitor cocktail (Roche) and phosphatase inhibitor cocktail (Roche). To assess the effects of reducing transcription, whole cell lysates were prepared from DMSO or CDK8i treated cells. 1x10^6^ cells were plated in 10 cm dishes and treated with 1 μM MSC2530818 (CDK8i) for 72 hours before they were collected and processed.

For detection of histone proteins and their modifications, total core histones were extracted from actively growing 5-10x10^6^ cells using the EpiQuik total histone extraction kit as per the manufacturer’s instructions. For detection of acetylated histones, panobinostat (10 μM) was included in each buffer during histone extraction.

Western blot analyses were performed on either whole cell lysates (30-40 mg) or total histone lysates (3-5 mg) that were separated on 4-15% SDS-PAGE and transferred to PVDF membrane. Membrane blocking and primary antibody incubations were performed in 5% non-fat milk dissolved in Tris-buffered saline, except for detection of phosphorylated proteins where BSA was used. Primary antibody incubations were performed overnight at recommended dilutions for each antibody before incubating with HRP-labeled secondary antibody and detection with ECL reagent. Primary antibodies are listed in the Supplementary Table S2.

### Hypertranscription in single nuclear (sn)-RNA-seq data from DMG patient tumor sample

To calculate the transcriptional output in single cells, we used an established measure for total transcriptional volume from the software package ISnorm (*49*). Briefly, ISnorm constructs in silico spike-in genes that are consistently expressed across cells by first creating a distance matrix between genes based on their transcriptional similarity. Next, it uses HDBSCAN to cluster genes that have mutually low distances, indicating their stability across cells. The relative abundance of these genes at a cell level is used to calculate the size factor, which is centered around 1 and is an estimate of the transcriptional volume of a given cell. To evaluate hypertranscription in DMG primary tumor samples, we applied ISnorm on published single-cell RNA-sequencing data from Jessa et al. (*19*), restricted to patients profiled with snRNA-seq. Cell doublets in the snRNA-seq datasets were identified and removed using DoubletFinder (*50*). ISnorm was then applied with a detection rate of 0.5 and otherwise default parameters. To statistically quantify effects of the H3K27M mutation on transcriptional output in tumor cells, we used a linear mixed-effects model (LMM), in which we additionally accounted for patient-level variability, as well as global differences in transcriptional output between tumor vs normal cells, or between H3K27M-mutated patients and patients who do not carry the mutation. Specifically, our model fits ISNORM RNA output scores using the following terms: a fixed effect that captures whether a cell is a tumor or normal cell, a fixed effect that captures if a patient’s tumor is H3 wild-type or H3K27M mutated, an interaction term to quantify potential combined effects between cell type and mutation status, and a patient-level random effect. Formally, we estimated:

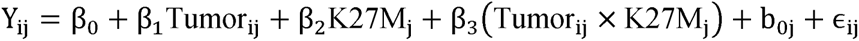

where Y_ij_ is the ISNORM-estimated RNA output for cell i in patient j, Tumor_ij_ is a binary indicator for malignant versus normal cells, H3K27M_ij_ denotes the histone mutation status of the patient, b_0j_ is a patient-specific random intercept, and □_ij_ is the residual error.

Patients were classified as H3K27M mutant if their tumor contained a H3K27M mutation (H3.3 or H3.1) and were classified as wild-type otherwise. All cells within each patient sample were classified as malignant or normal according to their annotation in Jessa et al.(*19*) The LMM model was fitted using Restricted Maximum Likelihood (REML) via the Python statsmodels package. The effects of H3K27M mutation on transcriptional output in tumor cells were thus quantified by the coefficient of the interaction term. Additional validation models were performed, including (1) replacing ISNORM RNA output with raw total counts, and (2) restricting analysis to malignant OPC and normal immune cells (using published cell annotations (*19*)), as these were the most abundant malignant and normal cell types, respectively.

### DNA fiber assay

To document replication perturbations, replicating DNA was labeled and analyzed by DNA fiber assays. Briefly, cells were pulsed with 100 mM of CldU (Sigma-Aldrich, C-6891) for 30 minutes, washed extensively, and again pulsed with 200 μM of IdU (Sigma-Aldrich, I-7125) for 30 minutes. To document fork stability and fork restart, HU (4 mM) was included as indicated by the schematics (Figure S3). At the end of the treatments, cells were immediately harvested, and 2.5x10^5^ cells were embedded in agarose plugs and treated with proteinase K solution (1000 mg of Proteinase K in lysis buffer {10 mM Tris-HCl [pH8.0], 100 mM EDTA [pH 8.0] and 1% N-lauroyl-sarcosine}) for 16 hours at 50°C. On the following day, plugs were washed extensively, melted in MES buffer at 68°C for 20 minutes, and digested with β-agarase at 42°C for 16 hours before DNA solutions were combed on coated coverslips (Genomic Vision) using the FiberComb Molecular Combing System. Combed coverslips were dehydrated for 2 hours at 60°C, denatured in freshly prepared 0.5 M NaOH for exactly 8 minutes at room temperature, washed, and again dehydrated sequentially in 70%, 90%, and 100% ethanol (3 minutes each).

Next, immunodetection of the replication tracks was performed using primary antibodies diluted in 1% BSA [Rat anti-BrdU to detect CldU (Abcam, ab6326); mouse anti-BrdU to detect IdU (BD Biosciences, 347580)] overnight at 4°C followed by washing and incubation with secondary antibodies-goat anti-rat IgG-Cy5 (Abcam, ab6565) and goat anti-mouse IgG-Cy3 (Abcam, ab6946) for 45 minutes at 37°C. To visualize single-stranded DNA (ssDNA), coverslips were also stained with mouse anti-ssDNA (DSHB University of Iowa) for 75 minutes at 37°C and detected with goat anti-mouse BV480 (BD Biosciences, 564877) for 45 minutes at 37°C. During immunodetection, all washes (repeated three times) were performed with 1X PBS containing 0.05% Tween-20 for 3 minutes. Finally, coverslips were dehydrated as before sequentially in 70%, 90%, and 100% ethanol, air-dried, mounted on slides, and scanned using the Fiber Vision automated scanner (Genomic Vision). The total length of IdU and the CldU labeled tracks were measured for >100 DNA fibers using CaseViewer 2.4, and to determine fork speed (kbp/min), total length was divided by 60 (total time for CldU and IdU incorporation) and multiplied by a factor of 2.59 (*51*) (Jackson and Pombo, 1998). To measure inter-origin distance, two consecutive CldU-IdU labeled tracks on the same DNA strand were identified and the distance between the centers of the two tracks was measured. To quantitate fork collisions, number of colliding forks was represented as the percentage of total forks.

### Quantification of chromatin-bound RPA2 and phosphorylation of CHK1^S317^

To quantify replication stress and ATR activation, we used flow cytometry-based approaches to measure chromatin-bound RPA2 and phosphorylation of CHK1 at ser317, respectively. All experiments were performed in triplicate. EdU (10 mM) was added to actively growing cells (∼60% confluency) for 45 minutes before cells were harvested for further processing. For chromatin-bound RPA2 detection, cells were pre-extracted (pre-extraction buffer:10 mM HEPES-KOH [pH 7.4], 300 mM sucrose, 100 mM NaCl, 3 mM MgCl_2_ and 0.3% Triton-X100) on ice for 10 minutes, fixed in 4% paraformaldehyde for 15 minutes, and resuspended in 1X saponin buffer for click-it chemistry to detect EdU. To detect CHK1^S317^ phosphorylation, cells were simultaneously fixed and permeabilized using BD Cytofix/Cytoperm buffer (BD Biosciences, 554714) for 30 minutes on ice and resuspended in 1X saponin buffer to detect EdU by click-it chemistry. Click-it chemistry was performed as described in the cell cycle analysis section (kit from ApexBio, K1179). Cells were washed twice with 1X PBS containing 0.5 mg/ml BSA, incubated in blocking buffer (1X PBS containing 3% goat serum, 0.1% Triton X-100, 1 mM EDTA and 1 mg/mL BSA) for 1 hour before incubation with primary antibody (anti-RPA2 at 1:200, ab2175, Abcam Inc.; anti-phospho-CHK1^S317^ at 1:100, 12302S, Cell Signaling Technology) in blocking buffer overnight at 4°C. After overnight incubations, cells were washed, incubated with fluorophore-conjugated secondary antibody solution for 1 hour, washed, and resuspended in 1X PBS containing 0.5 mg/ml BSA and Hoescht 33342. Cells were analyzed by flow cytometry using CYTOFLEX S and quantified by FlowJo software.

### Cell treatment and drug sensitivity assays

To measure ATRi sensitivity by colony formation assays, 3x10^3^ cells were plated in 10 cm plates, allowed to adhere for 16-18 hours and alnodesertib (ATRi) was added as indicated in the figures. Cells were allowed to grow for 9 days to form colonies. To visualize the colonies, colonies were fixed (10% methanol and 10% acetic acid) for 15 minutes and then stained with 0.25% crystal violet stain (resuspended in 10% ethanol) for 1 hour. Plates were thoroughly washed with water and colonies were counted. To measure drug sensitivity by CellTiter-Glo assays, isogenic SF188-H3.3^WT^, SF188-H3.3^K27M^, immortalized human astrocytes (IHA), and retinal epithelial cell (RPE-1) cells (250 cells/well), isogenic murine DMG cell lines (5000 cells/well), and patient-derived DMG cells (500 cells/well) were plated in triplicate or sextuplicate in 96-well plates. Cells were treated as indicated in the figures and cell viability was assessed by CellTiter-Glo assay after 6 days. To determine and visualize the synergy or antagonism relationship, Combenefit software was used (*33*).

### Proximity ligation assay (PLA) and R-loop detection

To detect transcription-replication conflict events, PLAs were performed with the indicated antibodies according to the manufacturer’s instructions (Sigma-Aldrich, DUO92101). For each assay, one of the antibodies was raised in mouse while the other was raised in rabbit. Cells were plated in fresh media overnight, and next day EdU (10 mM) was added for 45 minutes before cells were processed for PLA. Cells were pre-extracted in 1X PBS containing 0.5% NP-40 for 5 minutes on ice, fixed with 4% paraformaldehyde for 15 minutes and re-permeabilized in 1X PBS containing 0.5% Triton-X100 for 10 minutes. Next, cells were incubated in 1X PBS containing 5 mM EDTA and 100 mg of RNASE A for 30 minutes at 37°C, washed, and EdU click-it chemistry reaction was performed as described for Cell cycle analysis (ApexBio, K1179). Cells were washed and blocked in blocking buffer. For DMG cell lines, cells were cytospun on glass slides in blocking buffer. After 1 hour of blocking, cells were incubated overnight at 4°C with primary antibodies that were diluted (1:1000 for each antibody) in antibody diluent. Next day, cells were washed with PLA wash buffer A and incubated with anti-mouse and anti-rabbit probes for 1 hour at 37°C. Unhybridized probes were washed and followed by ligation for 30 minutes at 37°C and amplification for 1.5 hours at 37°C before the cells were mounted in anti-fade mounting media with DAPI and stored at -20°C until imaging. Cells were imaged on a Zeiss microscope at 63X magnification, and Cell Profiler software (*52*) was used to quantify PLA dots in nuclei and identify S phase cells (EdU positive).

To detect R-loops by dot blots, genomic DNA was isolated from ∼4x10^6^ actively growing cells as described (*53*). Cells were lysed in 600 ml cold lysis buffer (5 mM PIPES [pH8.0], 80 mM KCl, 0.5% NP-40) on ice for 10 minutes before nuclei were collected by centrifugation and resuspended in 800 ml cold nuclear lysis buffer (25 mM Tris-HCl [pH8.0], 1% SDS, 5 mM EDTA) for 5 minutes on ice. Next, nuclei were treated with 120 mg of Proteinase K at 55°C for 16 hours. Next day, equal volume of elution buffer (10 mM Tris-HCl [pH 8.5]) was added before genomic DNA was purified by phenol: chloroform extraction and ethanol precipitation. Glycogen was used as a carrier during ethanol precipitation. Genomic DNA (2 mg) was treated with RNASE T1 (400 Units) and RNASE III (1.5 Units) in 1X RNASE III buffer for 20 minutes at 37°C. Parallel reactions were further treated with RNASE H1 (5 Units) for 20 minutes at 37°C. Next, 250 ng and 500 ng equivalent reaction mixtures were spotted on two positively charged nylon membranes, one for S9.6 antibody and one for dsDNA antibody, as indicated in the figure. Once the membrane was dry, they were UV cross-linked using auto settings on the Stratalinker UV1800. Next, membranes were blocked, incubated with primary antibody, and detected as described for western blotting.

To detect R-loops using chromatin-bound V5-tagged RNASEH1^D210N^, immunofluorescence staining was performed with anti-V5 antibody on SF188-H3.3^WT^/V5-RNASEH1^D210N^ and SF188-H3.3^K27M^/V5-RNASEH1^D210N^ isogenic human glioma cells. Cells were plated on coverslips overnight, pre-extracted in 1X PBS containing 0.5% Triton-X100 on ice for 10 minutes and fixed in 4% paraformaldehyde for 15 minutes. Cells were blocked in 1X PBS containing 3% goat serum, 0.1% Triton X-100, 1mM EDTA, and 1 mg/mL BSA for 1 hour and incubated with primary antibody (V5 at 1:200, #903802, Biolegend) in blocking buffer overnight at 4°C. After overnight incubations, slides were washed, and the V5 signal was detected with AlexaFlour-594-conjugated secondary antibody solution for 1 hour. Nuclei were counterstained with DAPI, imaged at 63X magnification using the Zeiss microscope, and V5 staining intensity per nucleus was determined using CellProfiler software (*52*) for at least 100 cells.

### γ-H2AX, micronuclei, and apoptosis detection assays

Cells were plated in fresh media overnight before they were treated with either alnodesertib (500 nM or 1000 nM), radiation (1.5 or 2 Gy), or a combination thereof for assays described below.

To measure DNA damage, 2x10^5^ cells were plated in triplicate per condition, and γ-H2AX assay was performed as previously described (*54*) after 48 hours of treatment. Nuclei were stained with Hoechst 33342, data were collected on CytoflexS and analyzed by FlowJo software.

For the detection of micronuclei, 1x10^5^ cells were plated on coverslips and fixed 24 hours after treatment with 4% paraformaldehyde in 1X PBS for 15 minutes. Cells were washed three times in 1X PBS containing 0.5% Triton-X100 for 3 minutes each and mounted in DAPI-containing flouromount. Slides were imaged at 63X magnification with a Zeiss microscope, and images were scored manually for the presence of micronuclei in at least 100 cells.

To quantify apoptosis, 2.5x10^3^ cells were plated per well in 96 well plate in triplicate and Caspase-Glo 3/7 activity assays were performed 72 hours after treatment. Luminescence was read using CLARIOstar® microplate reader after 1 hour of incubation with the reagent and represented relative to untreated controls.

### Orthotopic DMG tumors, *in vivo* survival experiments and pharmacodynamic markers

All animal experiments performed in this study were approved by the Dana-Farber Institutional Care and Use Committee (IACUC) and were conducted as per NIH guidelines for animal welfare. Animals were housed and cared for according to standard guidelines with free access to water and food. All experiments were performed on 5-6 weeks-old female NSG mice (NOD.Cg-Prkdcscid Il2rgtm1Wjl/SzJ; Taconic Laboratories). Mice underwent intracranial surgeries for implantation of DIPG1 or SU-DIPG13-P* to establish orthotopic DMG tumors, randomly distributed among treatment groups, and treated as indicated. Mice were euthanized as they developed neurological symptoms or if they lost 15% of their body weight.

To establish orthotopic DMG xenografts, animals were injected intraperitoneally with the analgesic buprenorphine 0.05 mg/Kg, then anesthetized with isoflurane 2–3% mixed with medical air and placed on a stereotactic device. Next, a small incision and a small hole was made with a 25-gauge needle and 2 ul of 1X PBS containing 1x10^5^ DIPG1-luc or SU-DIPG13-P*-luc (cells expressing luciferase gene) cells were injected stereotactically into the right pons (stereotactic coordinates zeroed on bregma: -1.5 mm X(ML), -5.5 mm Y(AP) and -5.0 mm Z(DV)) at rate of 1μl/min with an infusion pump before the incision was sutured. Animal health was assessed daily for signs of distress, neurological symptoms, and weight loss. Tumor growth was monitored weekly using the IVIS Spectrum *In Vivo* Imaging System (PerkinElmer), starting at day 7 post-cell injections as described previously (*43*). Quantification of bioluminescence signal was performed using the Living Image Software (PerkinElmer).

For monotherapy drug studies, mice were administered alnodesertib at MTD (maximum tolerated dose;100 mg/Kg) or placebo (vehicle for alnodesertib resuspension: 5% Tween-80 and 0.5% methylcellulose in sterile water) by oral gavage, twice daily for 3 consecutive days, followed by 4 days drug-free. Treatment of mice was started 8-9 days after intracranial surgeries and mice were treated with alnodesertib for 8 weeks. For MALDI-MSI studies, alnodesertib was administered twice daily for 3 days at 100 mg/Kg before intracranial DIPG1-luc tumor bearing mice were euthanized and brains were collected. For studies, combining alnodesertib and radiation, tumor-bearing mice were randomized to one of the following treatment groups: (i) control; (ii) radiation (^137^Cs source); (iii) alnodesertib; (iv) alnodesertib + radiation. Alnodesertib was administered at half-MTD (50 mg/Kg) twice daily for 3 consecutive days followed by 4 days drug-free on a 7-day cycle. Radiation was administered at 0.5 Gy to the head, in between the two daily doses of alnodesertib for a total of 3 doses on a 7-day cycle. Radiation was administered for 2 weeks (6 x 0.5 Gy) and alnodesertib was given for 8 weeks.

For pharmacodynamic marker assessments, formalin-fixed brain tissues of randomly selected mice belonging to either vehicle or alnodesertib treatment groups in the efficacy study were paraffin-embedded, sectioned, immunostained, and quantified using Halo Indica services provided by iHisto (https://www.ihisto.io/).

### Tumor and tissue analysis by MALDI mass spectrometry imaging and microscopy

Mouse brains were dissected, snap-frozen, and stored at -80°C. Sagittal cryosections of 10 μm thickness were collected from each mouse brain and thaw-mounted onto indium-tin-oxide (ITO) slides for MALDI MSI. Serial sections were brush-mounted on glass microscopy slides for hematoxylin and eosin (H&E) staining. H&E-stained sections were imaged using a 10X objective on a Zeiss Observer Z.1. For drug quantification, alnodesertib was dissolved in DMSO and spiked into healthy mouse control brain homogenate at various concentrations (0.00, 0.10, 0.50, 1.00, 5.00, and 8.00 µM) to create a tissue mimetic model. Spiked tissue homogenates were dispensed into gelatin tissue microarray molds and cryosectioned onto ITO slides at 10 µm thickness alongside the experimental samples. Prior to data acquisition, the ITO slide containing experimental and tissue mimetic sections was placed in a desiccator for 10 minutes, then uniformly coated with a matrix solution of 2,5-dihydroxybenzoic acid (160 mg/mL) in 70:30 methanol:0.1% trifluoroacetic acid with 1% DMSO. MALDI matrix solution was applied in a crisscross pattern over a two-pass cycle using an M5 Sprayer (HTX Technologies, Chapel Hill, NC) with the following spray parameters: 0.18 mL/minute flow rate, 75°C spray nozzle temperature, 1200 mm/minute nozzle velocity, 40 mm nozzle height, 2 mm track spacing, and 10 psi nitrogen gas pressure.

Alnodesertib distribution was mapped using a timsTOF flex mass spectrometer (Bruker Daltonics, Billerica, MA) in positive ion mode with a selected reaction monitoring method scanning the range of *m/z* 100 to *m/z* 2000. Collision energy was set to 50 eV, and the transition *m/z* 389.18 [M+H] → *m/z* 333.11 was monitored using a 3 *m/z* isolation window. Transfer funnel, collision cell, quadrupole, and focus pre-TOF method parameters were optimized by direct infusion of alnodesertib solution through the ESI source. MALDI source parameters were set to 50 µm raster size, 1000 laser shots per pixel, 5000 Hz laser repetition frequency, and 74% laser power (arbitrary units). After alnodesertib imaging, heme b distribution was acquired in positive ion mode using a full-scan detection method across the same tissue raster spots. The scanning range was set from *m/z* 100 to *m/z* 1200, and the MALDI source parameters were set to 200 laser shots per pixel, 10000 Hz laser repetition frequency, and 70% laser power (arbitrary units). The instrument was calibrated with Agilent tune mix solution (Agilent Technologies, Santa Clara, CA) using the ESI source prior to both imaging acquisitions.

Alnodesertib and heme b distributions were visualized and analyzed in SCiLS Lab (Bruker Daltonics, Billerica, MA). Tumor and non-tumor regions were defined by co-registration of H&E images with the MSI regions within the software. Unnormalized intensity values of the drug fragment peak were exported from tumor, non-tumor, and tissue mimetic regions. OriginPro software (OriginLab Corp., Northampton, MA) was used to plot tissue mimetic concentrations against the detected intensities, creating a linear calibration curve with instrumental weighting used to calculate the tumor and non-tumor drug concentrations from their measured intensity values. The limit of detection and limit of quantification were defined as 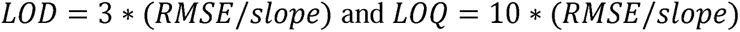, respectively.

### *In situ* replication stress detection and quantification

To quantify replication stress *in situ*, dSTRIDE-pRPA assay was performed. Briefly, *in situ* detection of double-strand breaks (DSBs) associated with phosphorylated RPA (pRPA) was performed using the dSTRIDE-pRPA assay on brain tissue sections of mice harboring intracranial DMG tumors. The assay utilizes the principles of STRIDE previously described in Kordon et al.(*55*). Briefly, modified nucleotide analogs (BrdUTP) were enzymatically incorporated at DSB ends and then detected using appropriate primary antibodies alongside detection of phosphorylated RPA with anti-pRPA32 antibodies (Abcam, ab211877). Next, slides were incubated with secondary antibodies that are conjugated to oligonucleotides followed by ligation to form a circular DNA template. Then, rolling circle reaction was performed to generate the amplicon, and finally, to detect the amplicon, short fluorescently labelled oligonucleotides were hybridized and mounted in DAPI-containing mounting media. Slides were imaged using a Zeiss Celldiscoverer 7 LSM 900 confocal microscope. At least seven fields of view (FOVs) were acquired per tissue sample, focusing on tumor-rich regions. Images were captured as 3D confocal stacks, nuclei were segmented based on DAPI signal to create 3D masks, and dSTRIDE-pRPA foci within the nuclei masks were identified and quantified using 3D local maxima detection method.

### Statistical analysis

Data analysis and visualization were performed with PRISM 9 (GraphPad software). Unlessindicated, all data were represented as mean ± SEM and plotted using PRISM software. Area under the curve (AUC) for drug responses was calculated using the PRISM software. For two group comparisons, the t-test was used. Whenever multiple groups were compared, one-way ANOVA was used, and Tukey test was performed to correct for multiple comparisons in PRISM. For datasets that failed normality test, we performed nonparametric Kruskal-Wallis testing with Dunn’s test to correct for multiple comparisons in PRISM.

For the statistical analysis of transcriptional output from snRNA-seq datasets of DMG primary tumor tissues, a linear mixed-effects model (LMM) was employed as detailed in the methods.

For dSTRIDE-pRPA assays, statistical analysis was performed using generalized linear mixed models (GLMMs) with a negative binomial distribution to account for overdispersion and the hierarchical nature of the dataset (cells nested within FOVs and samples). Treatment was defined as a fixed effect, while nested random intercepts were used to partition variance across all experimental levels. Statistical significance was determined via Holm-adjusted pairwise contrasts comparing treatment groups to the vehicle control.

Asterisks indicate statistically significant values (*, p<0.05; **, p<0.01; ***, p<0.001; ****, p<0.0001).

## Supplementary Materials

Materials and Methods

Fig S1 to S7

Table S1 and S2

## Acknowledgement

We are grateful to Dr. David Allis for the expression plasmids of histone H3.3, to Dr. Timothy Phoenix for the mouse DMG cell lines, and to Drs Esther Hulleman and Michelle Monje for human DMG cell lines. We also thank Dr. Brendan Price for supervising the generation of SF188 isogenic cell lines, Dr. Shrabasti Roychowdhury for helpful discussions, Dr. Ke Cong and Vaishnavi Anand for help with establishing DNA fiber assays, Tori Donovan for intracranial DMG xenograft establishment, and Dr. Andrey Aristov for initial assessment of tumor tissue staining. The research was supported in part by the William M. Wood Foundation (D.H-K), UCSF-DFCI donor grant GR0130774-S01(D.H-K and S.M), Research Grants-NCI SPORE grant 5P50CA165962 (D.H-K), PAIRS grant 1R01CA284565 (D.H-K and D.C), U54 grant 5U54CA274516 (D.H-K, F.M, S.M), Innovations Research Fund of Dana-Farber Cancer Institute (D.H-K), Daniel E. Ponton Fund for the Neurosciences (NYRA), and the National Brain Tumor Society DDR Consortium (NYRA).

## Author contributions

Conceptualization, S.P, N.Y.R.A, F.M and D.H-K; Methodology development and support, S.P, T.M, H.W, S.M, T.K, A.N, K.K, J.G, C.J.G, J.B.M, H.R, G.S, K.B, N.Y.R.A, O.W, M.B and D.H-K; Investigation, S.P, H.W, J.G, C.J.G, A.N, T.K, K.K, T.M, O.W, M.B; Validation, S.P, H.W, N.Y.R.A, J.G, C.J.G and D.H-K; Formal analysis and Data review, S.P, H.W, N.Y.R.A, T.M, K.K, J.B.M, H.R, J.G, C.J.G, O.W, M.B, D.C; Visualization, S.P, H.W, N.Y.R.A, J.G, C.J.G, O.W, M.B; Writing – Original Draft, S.P, C.S, D.H-K; Writing – Review & Editing, All authors; Initiating discussions and collaboration, S.J.B, G.S; Funding Acquisition, D.H-K, S.M, D.C, F.M and N.Y.R.A; Resources, S.P, T.M, H.W, N.Y.R.A, and D.H-K; Project administration, D.H-K; Supervision, D.H-K., S.P, D.C, S.M, F.M and N.Y.R.A.

## Data and materials availability

DMG primary tumor tissue single cell nuclear RNA sequencing data used for hypertranscription analysis in this study has been generated and published by Jessa et al.2022 and is deposited as described in the publication (*19*). All other data are available in the main text or the supplementary materials. All raw data generated for the study are available upon request.

