## Supplementary File for "H3K27M-driven hypertranscription leads to a new targetable dependency in diffuse midline gliomas"

For detection of histone proteins and their modifications, total core histones were extracted from actively growing 5-10x$10^6$ cells using the EpiQuik total histone extraction kit as per the manufacturer's instructions. For detection of acetylated histones, panobinostat (10 mM) was included in each buffer during histone extraction.

Supplementary Figures and Legends

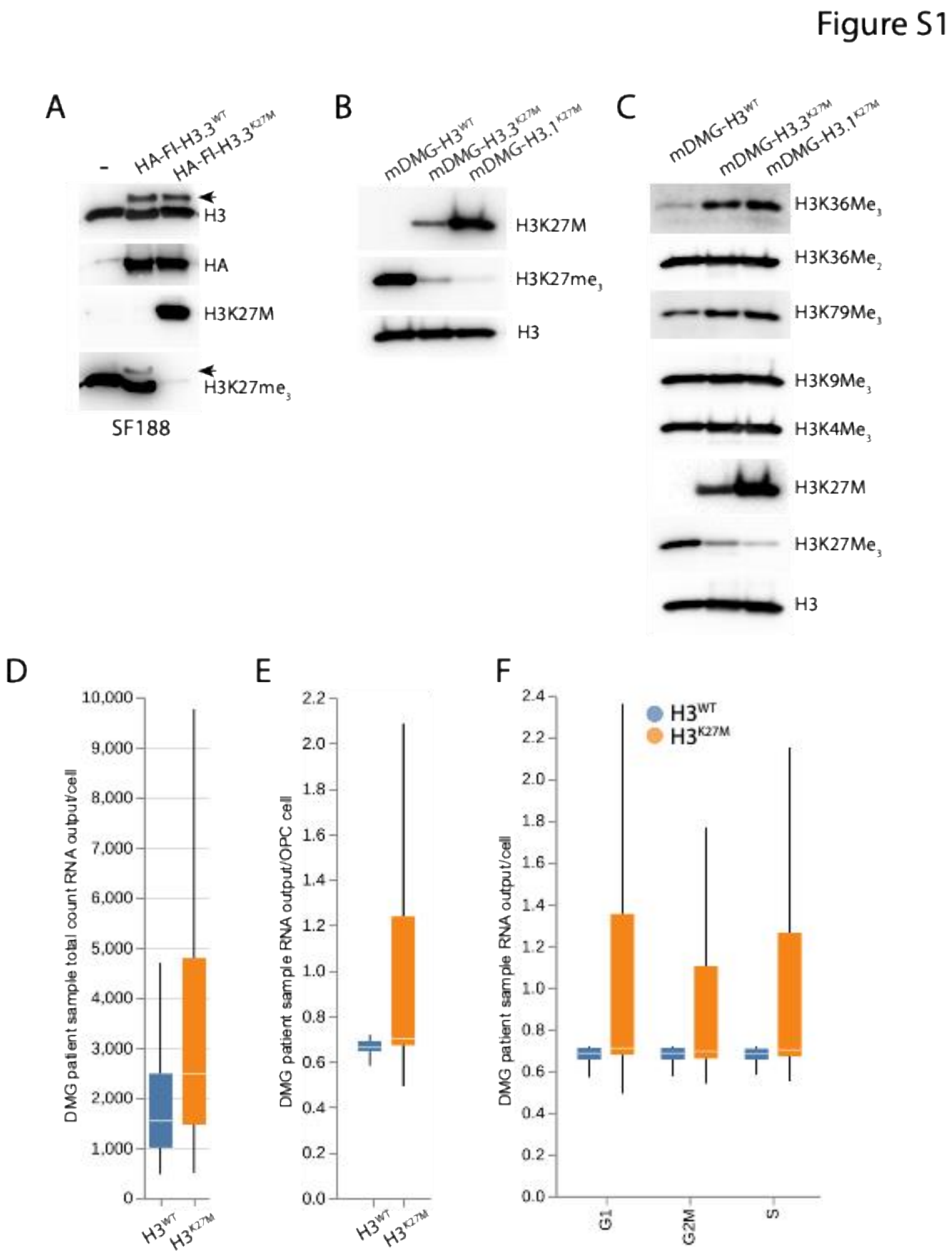

**Fig. S1: Characterization of isogenic human and mouse cells and primary DMG specimens.**

**(A)** Generation of isogenic pairs expressing wild-type histone H3.3 (HA-FLAG-$H3.3^{WT}$) or H3K27M-mutated H3.3 (HA-FLAG-$H3.3^{K27M}$) in the pediatric high-grade glioma cell line SF188. Western blot analysis shows expression of FLAG- and HA-tagged H3.3, H3K27M, and loss of $H3K27me_3$ in H3K27M-expressing cells. The parental SF188 is included for comparison. **(B)** Western blot analyses of cells from isogenic murine DMG models confirm expression of H3K27M and loss of $H3K27me_3$. **(C)** Western blot analyses of transcription-associated methylation of total histones in murine isogenic DMG cells. **(D-F**) Box plots of RNA output (transcriptional volume) of single cells sequenced from DMG primary tumor specimens based on (D) total RNA-seq counts, (E) cells identified as OPCs and normal immune cells, and (F) cell cycle phase.

Figure S2

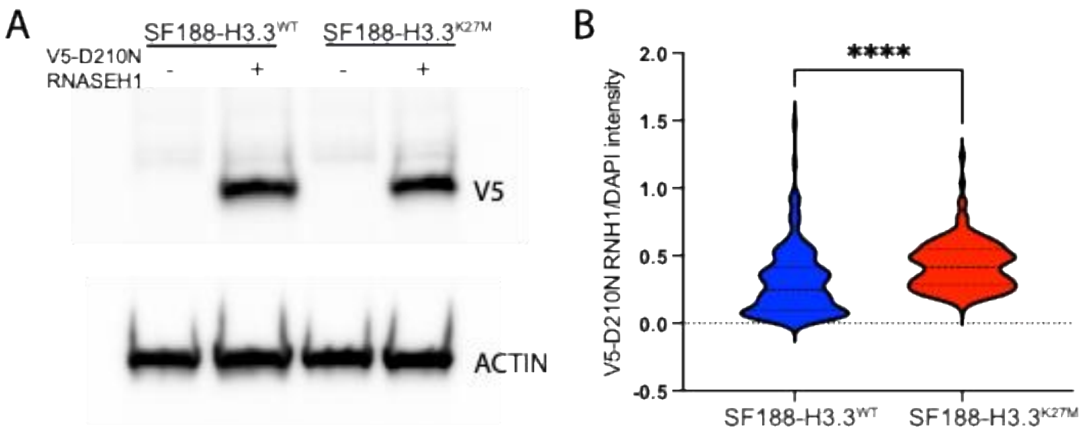

**Fig. S2: RNASEH1$^{D210N}$ mutant detects increased RNA-DNA hybrids in H3K27M cells.** **(A)** Isogenic SF188-H3.3$^{WT}$ and SF188-H3.3$^{K27M}$ cells were transduced to express V5 tagged-RNASEH1$^{D210N}$ mutant, which binds RNA-DNA hybrids, to detect levels of genomic R-loops. Western blots using anti-V5 antibody confirm expression of mutant RNASEH**1. (B)** Immunofluorescence signal from anti-V5 probing of at least 100 pre-extracted cells, quantified by CellProfiler software and represented as violin plots.

### Figure S3

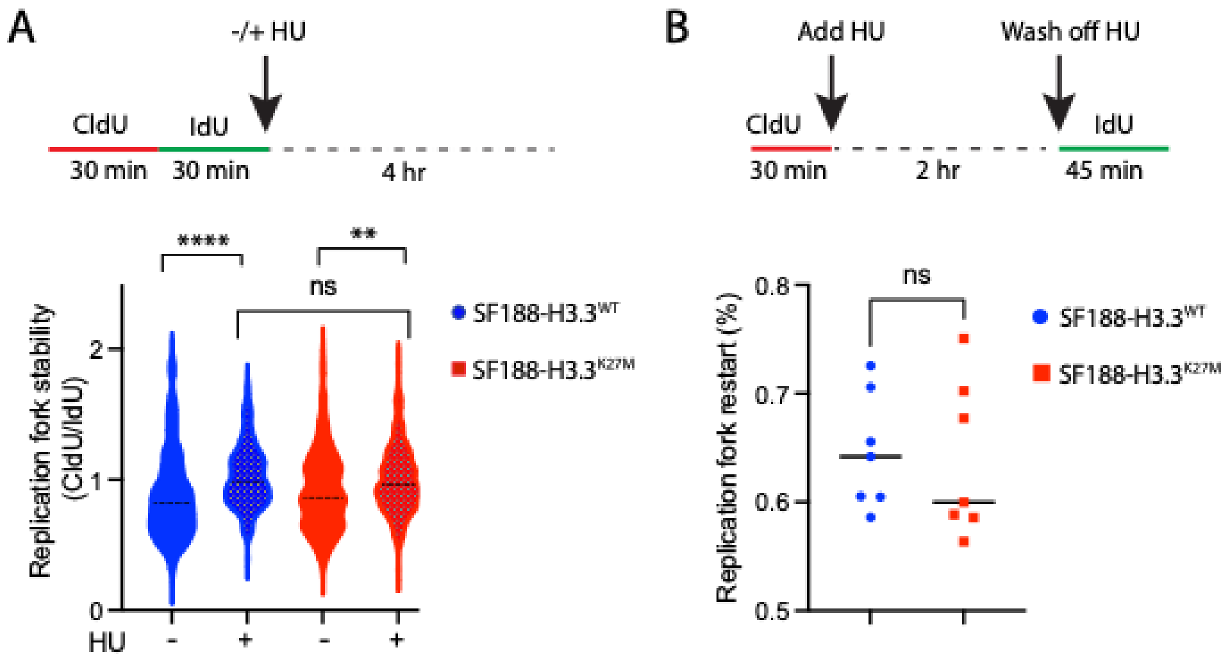

**Fig. S3: H3K27M does not impact replication fork stability or restart.**
**(A and B)** DNA fiber assays were performed in isogenic SF188-H3.3$^{WT}$ and SF188-H3.3$^{K27M}$ cells to measure (A) replication fork stability and (B) replication fork restart according to the schematics in upper panels. Replication fork restart is calculated as the number of forks with continuous red-green tracks relative to the total number of forks [red-green/(green only + red only + red-green); n=6 from 2 independent experiments; each data point represents non-overlapping regions with at least 100 replication tracks]. Replication fork stability is determined by calculating the CldU/IdU ratio (pooled data from two independent experiments and >100 tracks were quantified per experiment).

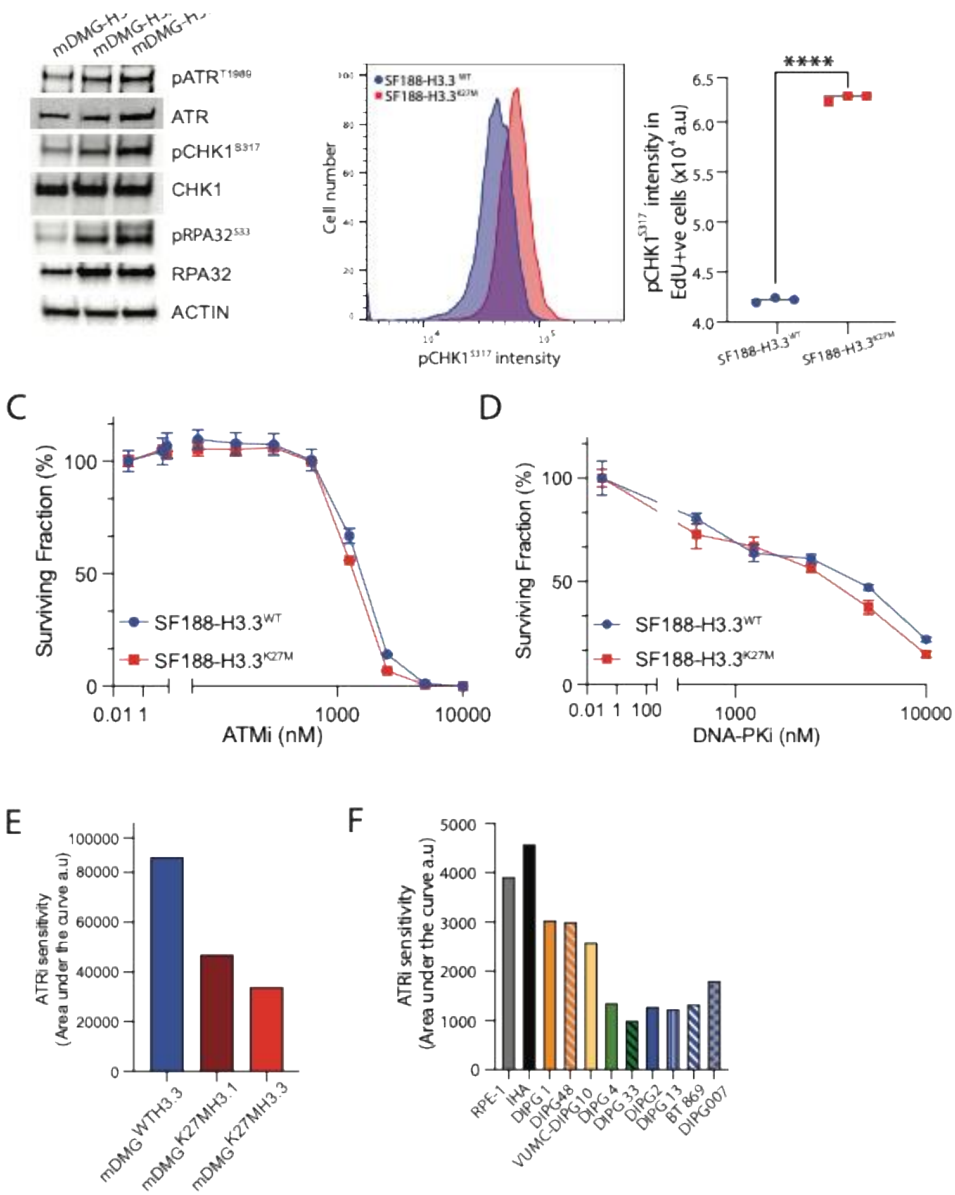

**Fig. S4: H3K27M drives ATR activation and sensitivity to ATR inhibition.**

**(A)** Western blot analyses of cells from isogenic murine DMG models confirm activation of ATR kinase in H3K27M-mutated cells. **(B)** Phosphorylation of CHK1 at S317 was measured by flow cytometry in isogenic SF188-H3.3$^{WT}$ and SF188-H3.3$^{K27M}$ cells. Representative histograms (left panel) and quantification (n=3; right panel) are shown. **(C and D)** Cell-titer Glo assays on SF188-H3.3$^{WT}$ and SF188-H3.3$^{K27M}$ cells using inhibitors of (C) ATM or (D) DNA-PK. **(E and F)** Area under the curve, shown by bar chart, for ATRi (alnodesertib) sensitivity shown in Figures 4F and 4G.

Figure S5

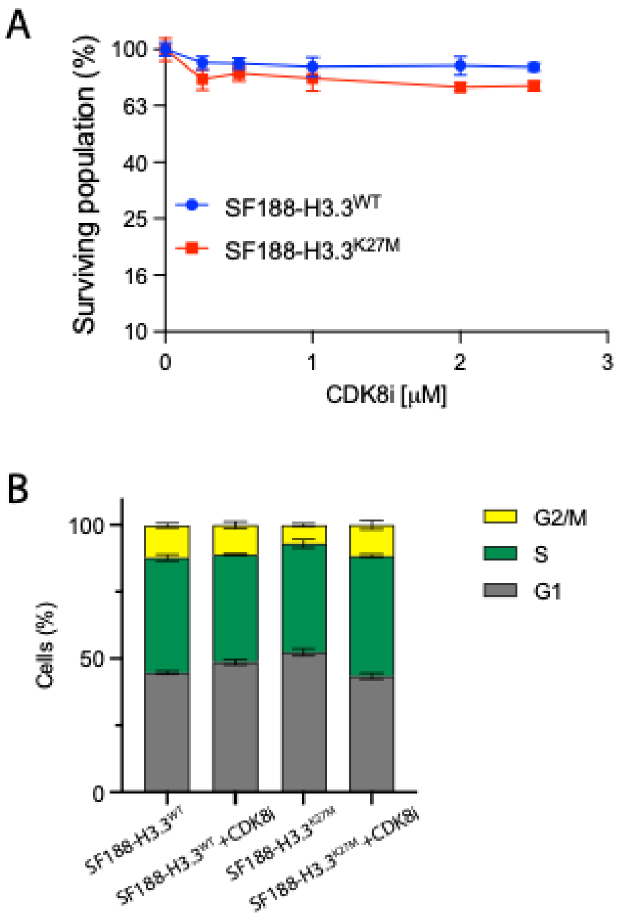

**Fig. S5: CDK8 inhibition has minimal effect on cell viability and cell cycle profile.**
**(A)** Isogenic SF188-H3.3$^{WT}$ and SF188-H3.3$^{K27M}$ cells were treated with CDK8i and cell viability was determined by CellTiter-Glo assays after 5 days (mean± SEM, n=5). **(B)** Cell cycle profile of isogenic SF188-H3.3$^{WT}$ and SF188-H3.3$^{K27M}$ cells with or without CDK8i treatment (1 mM, 72 hours; mean± SEM, n=3).

Figure S6

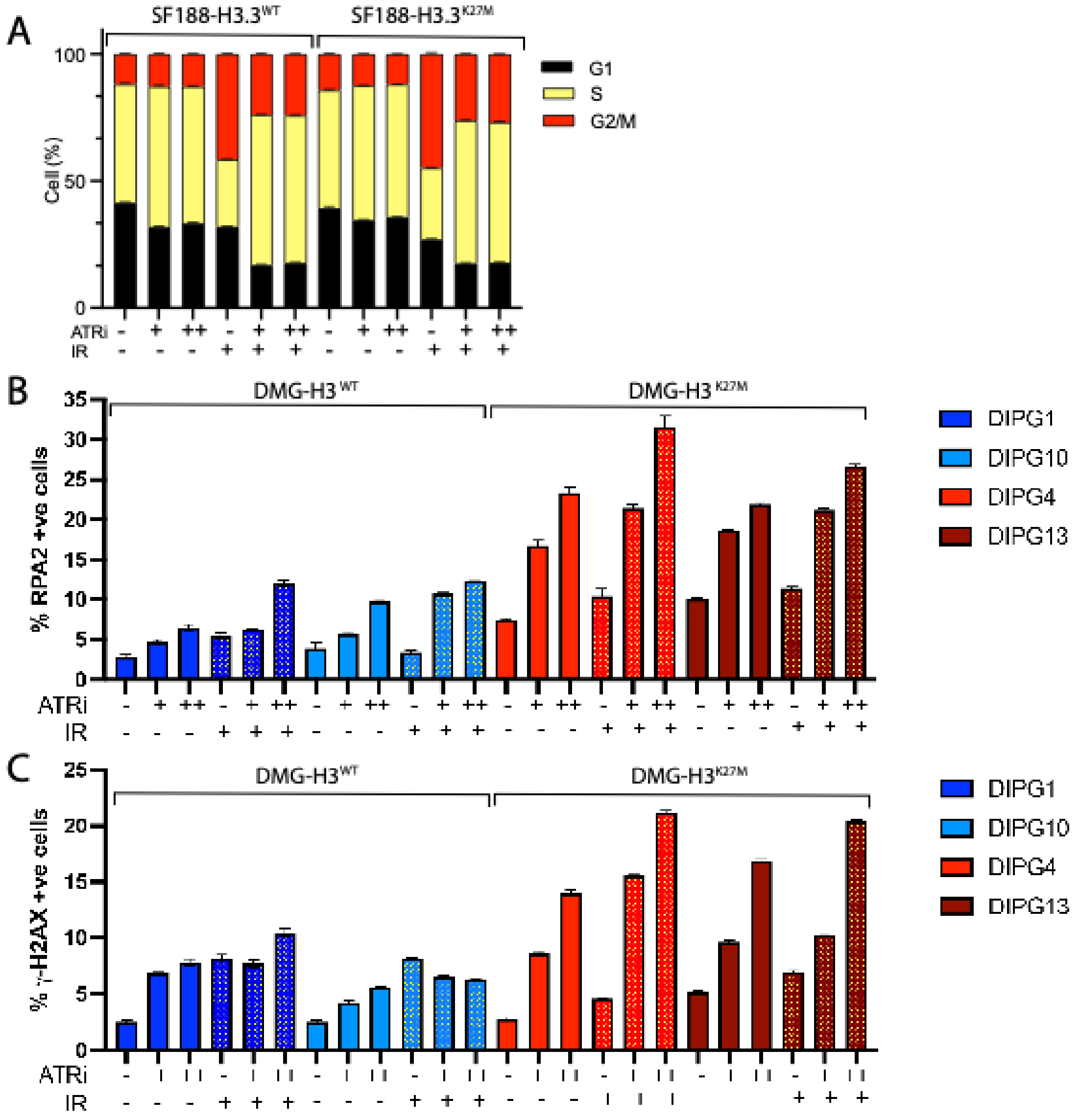

**Fig. S6: Radiation enhances replication stress and DNA damage in DMG cells, most prominent in H3K27M cells.**

**(A)** Cell cycle distribution of SF188-H3.3$^{WT}$ and SF188-H3.3$^{K27M}$ cells treated with ATRi and radiation as monotherapy or in combination, determined by flow cytometry (ATRi: 500 and 1000 nM; IR: 2 Gy; 24 hours; n=3). **(B)** Replication stress measured by chromatin-bound RPA (ATRi: 500 and 1000 nM; IR: 2 Gy; 24 hours; n=3), and **(C)** DNA damage measured by γ-H2AX (ATRi: 500 and 1000 nM; IR: 2 Gy; 48 hours; n=3) were assessed in H3 wild-type and H3K27M-mutated patient-derived DMG cell lines.

Figure S7

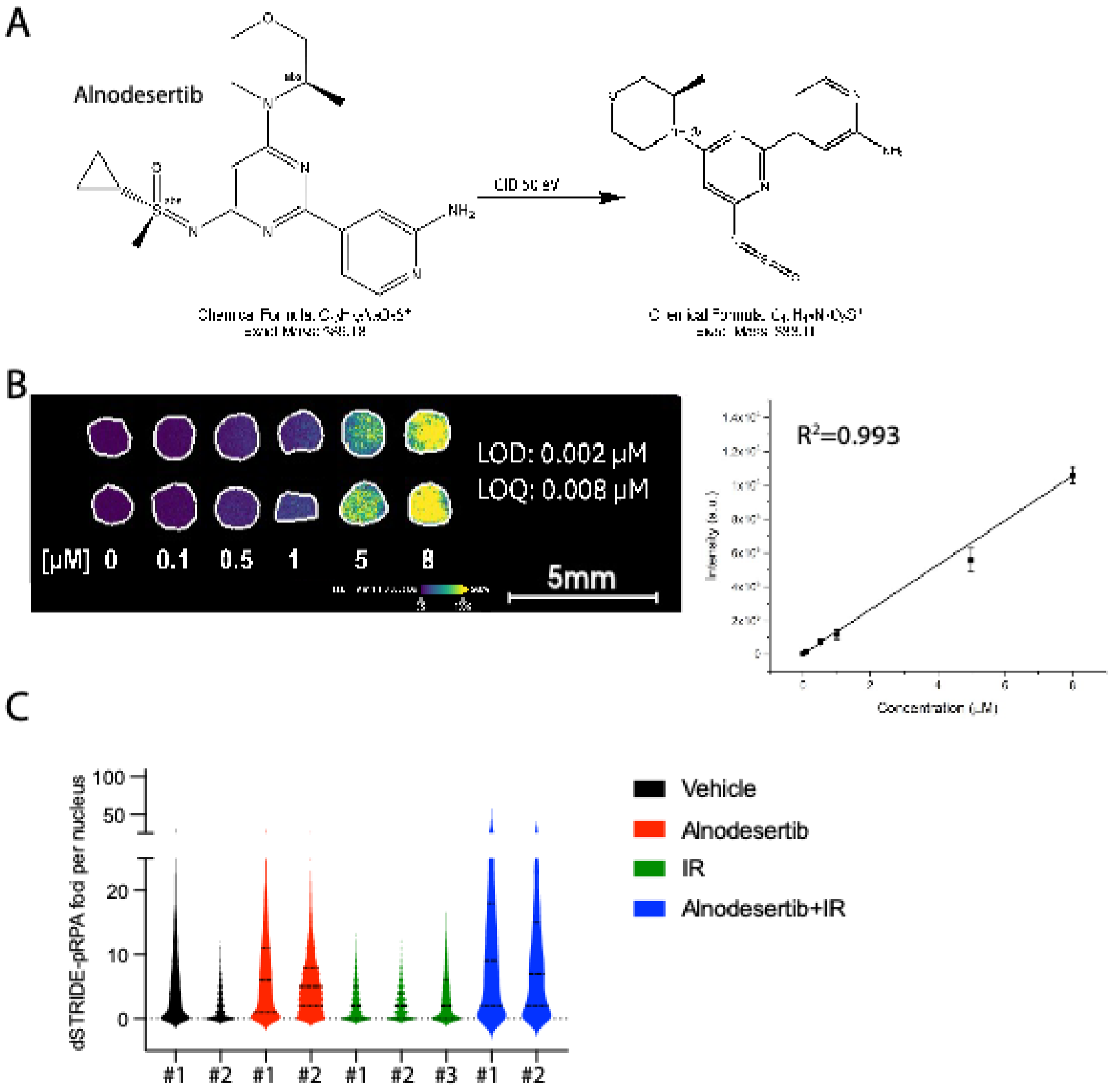

**Fig. S7: MALDI-MSI quantification of alnodesertib and quantification of colocalized DSB and pRPA in mouse brain tumor regions.**

**(A)** Chemical structure of alnodesertib and its fragmentation for MALDI-MSI detection. **(B)** Tissue mimetic analysis of alnodesertib with dose ranges of 0-8 mM used to generate calibration curve for MALDI-MSI quantification. The calibration curve has $R^2$=0.993. LOD is the limit of detection and LOQ is the limit of quantification. **(C)** Violin plots show the distribution of i*n situ* quantification of co-localized DSB and pRPA foci (dSTRIDE-pRPA) per nucleus in tumor sections from each mouse (n=2; except for IR arm with n=3) treated in the four-arm *in vivo* experiment in Figure 7F.

**Table S1**: **Statistical testing for RNA expression in DMG primary tumor tissues based on single cell RNA-seq data.**

| Cell Types | Target Measure | Cell Type Coefficient | Cell Type P Value | K27M Coefficient | K27M P value | Interaction Coefficient | Interaction P value |
|---|---|---|---|---|---|---|---|
| Malignant vs Normal | RNA Output | 0.033 | 0.564 | 0.529 | 0.237 | 0.149 | 0.011 |
| Malignant vs Normal | Total Count | 313.639 | 0.314 | 3255.976 | 0.209 | 864.508 | 0.007 |
| OPC vs Immune | RNA Output | 0.022 | 0.811 | 0.19 | 0.694 | 0.578 | 0 |

| **Summary Statistics** | **RNA output based (Malignant cell vs Normal cell)** | | | | | |
|---|---|---|---|---|---|---|
| Model: | MixedLM | | | | | |
| No. Observations: | 59679 | | | | | |
| No. Groups: | 16 | | | | | |
| Min. group size: | 404 | | | | | |
| Max. group size: | 15566 | | | | | |
| Mean group size: | 3729.9 | | | | | |
| Dependent Variable: | RNA Output | | | | | |
| Method: | REML | | | | | |
| Scale: | 0.6801 | | | | | |
| Log-Likelihood: | -73241.36 | | | | | |
| Converged: | Yes | | | | | |
| **Coefficients** | | | | | | |
| | **Coef.** | **Std.Err.** | **z** | **P>\|z\|** | **[0.025** | **0.975]** |
| Intercept | 0.743 | 0.404 | 1.839 | 0.066 | -0.049 | 1.535 |
| Tumor | 0.033 | 0.057 | 0.576 | 0.564 | -0.078 | 0.143 |
| K27M | 0.529 | 0.448 | 1.182 | 0.237 | -0.348 | 1.407 |
| Tumor x K27M | 0.149 | 0.059 | 2.546 | 0.011 | 0.034 | 0.264 |
| Group Var | 0.481 | 0.224 | | | | |

| **Summary Statistics** | **Total RNA count based (Malignant cell vs Normal cell)** |
|---|---|
| Model: | MixedLM |
| No. Observations: | 59679 |

| | | | | | | |
|---|---|---|---|---|---|---|
| No. Groups: | 16 | | | | | |
| Min. group size: | 404 | | | | | |
| Max. group size: | 15566 | | | | | |
| Mean group size: | 3729.9 | | | | | |
| Dependent Variable: | Total Count | | | | | |
| Method: | REML | | | | | |
| Scale: | 20609153 | | | | | |
| Log-Likelihood: | -587245.2 | | | | | |
| Converged: | Yes | | | | | |
| **Coefficients** | | | | | | |
| | **Coef.** | **Std.Err.** | **z** | **P>\|z\|** | **[0.025** | **0.975]** |
| Intercept | 1638.374 | 2337.093 | 0.701 | 0.483 | -2942.2 | 6218.99 |
| Tumor | 313.639 | 311.199 | 1.008 | 0.314 | -296.3 | 923.577 |
| K27M | 3255.976 | 2589.889 | 1.257 | 0.209 | -1820.1 | 8332.07 |
| Tumor x K27M | 864.508 | 323.099 | 2.676 | 0.007 | 231.246 | 1497.77 |
| Group Var | 16104214 | 1300.322 | | | | |

| **Summary Statistics** | **RNA output (OPC vs Immune cells)** | | | | | |
|---|---|---|---|---|---|---|
| Model: | MixedLM | | | | | |
| No. Observations: | 21660 | | | | | |
| No. Groups: | 16 | | | | | |
| Min. group size: | 77 | | | | | |
| Max. group size: | 4792 | | | | | |
| Mean group size: | 1353.8 | | | | | |
| Dependent Variable: | RNA Output | | | | | |
| Method: | REML | | | | | |
| Scale: | 0.5977 | | | | | |
| Log-Likelihood: | -25215.733 | | | | | |
| Converged: | Yes | | | | | |
| **Coefficients** | | | | | | |
| | **Coef.** | **Std.Err.** | **z** | **P>\|z\|** | **[0.025** | **0.975]** |

| | | | | | | |
|---|---|---|---|---|---|---|
| Intercept | 0.775 | 0.436 | 1.778 | 0.075 | -0.079 | 1.629 |
| OPC | 0.022 | 0.093 | 0.24 | 0.811 | -0.16 | 0.205 |
| K27M | 0.19 | 0.482 | 0.394 | 0.694 | -0.755 | 1.134 |
| OPC x K27M | 0.578 | 0.096 | 6.015 | 0 | 0.39 | 0.766 |
| Group Var | 0.542 | 0.264 | | | | |

**Table S2: Antibodies used in this study.**

| Antibodies | Source | Application | Identifier |
|---|---|---|---|
| Anti-ATR [2B5] | GeneTex | Western blotting | Cat# GTX70109, RRID:AB_368158 |
| Anti-ATR | Cell Signaling Technology | Western blotting | Cat# 2790 |
| Anti-ATR (phospho Thr1989) | GeneTex | Western blotting | Cat# GTX128145, RRID:AB_2687562 |
| Anti-RPA32/RPA2 antibody [9H8] | Abcam Inc. | Western blotting, flow cytometry | Cat# ab2175, RRID:AB_302873 |
| Anti-RPA32/RPA2 antibody | Cell Signaling Technology | Western blotting | Cat# 52448 |
| Anti-RPA32/RPA2 (phospho S33) | Abcam Inc. | Western blotting | Cat# ab211877, RRID:AB_2818947 |
| Anti-RPA32/RPA2 (phospho S33; E9N1T) | Cell Signaling Technology | Western blotting | Cat# 10148 |
| Anti-Chk1 (2G1D5) | Cell Signaling Technology | Western blotting | Cat# 2360S, RRID:AB_2080320 |
| Anti-Chk1 (phospho Ser317) | Cell Signaling Technology | Western blotting | Cat# 2344, RRID:AB_331488 |
| Anti-Chk1 (phospho Ser317) | Cell Signaling Technology | Flow cytometry | Cat# 12302S |
| Anti-KAP1 | Biolegend | Western blotting | Cat# 619301 |
| Anti-KAP1 (phospho S824) antibody [BL-246-7B5] | Abcam Inc. | Western blotting | Cat# ab243870 |
| Anti-ATM | Santa Cruz Biotechnology | Western blotting | Cat# sc377293 |
| Anti-ATM (phospho Ser1981) | Santa Cruz Biotechnology | Western blotting | Cat# sc-47739 |
| Anti-RNA POL2 (rabbit) | Abcam Inc. | Western blotting | Cat# ab26721 |
| Anti-RNA POL2 (phospho Ser 2) | Abcam Inc. | Western blotting, PLA | Cat# ab5095 |
| Anti-RNA POL2 (phospho Ser 5) | Abcam Inc. | Western blotting, PLA | Cat# ab5131 |
| Anti-RNA POL2 (mouse) | Millipore | PLA | Cat# 05-952-1-25UG |
| Anti-PCNA (rabbit) | Cell Signaling Technology | PLA | Cat# 13110S |
| Anti-PCNA (mouse) | Santa Cruz Biotechnology | PLA | Cat# sc-56 |
| Anti-dsDNA | Abcam Inc. | Dot blot | Cat# ab27156 |
| Anti-DNA-RNA hybrid (S9.6) | Kerafast | Dot blot | Cat # ENH001 |

| | | | |
|---|---|---|---|
| Anti-V5 tag | Biolegend | Western Blotting, Immunoflourescence staining | Cat# 903802 |
| Anti-β-Actin (8H10D10) | Cell Signaling Technology | Western blotting | Cat# 3700S, RRID:AB_2242334 |
| anti-γ-H2AX (FITC-conjugated) | Biolegend | Flow cytometry | Cat# 613404; RRID: AB_528919 |
| Anti-Histone H3 | Abcam Inc. | Western blotting | Cat# ab18521 |
| Anti-Histone H3 (mutated K27M; [EPR18340]) | Abcam Inc. | Western blotting | Cat# ab190631 |
| Anti-Tri-methyl-Histone H3 (Lys27) (C36B11) | Cell Signaling Technology | Western blotting | Cat#9733 |
| Anti-Acetyl-Histone H3 (Lys27) (D5E4) | Cell Signaling Technology | Western blotting | Cat# 8173 |
| Anti-Tri-methyl-Histone H3 (Lys36) | Cell Signaling Technology | Western blotting | Cat# 9763 |
| Anti-Di-methyl-Histone H3 (Lys36) | Active Motif | Western blotting | Cat# 39056; RRID: AB_2793207 |
| Anti-Tri-methyl-Histone H3 (Lys79) (E8B3M) | Cell Signaling Technology | Western blotting | Cat#74073 |
| Anti-Tri-methyl-Histone H3 (Lys 4) | Abcam Inc. | Western blotting | Cat# ab8580 |
| Anti-Acetyl-Histone H3 (Lys9) (C5B11) | Cell Signaling Technology | Western blotting | Cat# 9649 |
| Anti-Tri-methyl-Histone H3 (Lys 9) | Abcam Inc. | Western blotting | Cat# ab8898 |
| Anti-Mouse IgG - Horseradish Peroxidase linked whole antibody (sheep) | Cytiva | Western blotting | Cat# NA931V, RRID:AB_772210 |
| Anti-Rabbit IgG - Horseradish Peroxidase linked F(ab')2 fragment (donkey) | Cytiva | Western blotting | Cat# NA934V, RRID:AB_772206 |
| Anti-Mouse IgG - Alexa Flour (AF) 594 conjugated (donkey) | Invitrogen | Immunoflourescence staining, Flow cytometry | Cat# A21203 |
| Anti-Rabbit IgG - Alexa Flour (AF) 594 conjugated (donkey) | Invitrogen | Immunoflourescence staining, Flow cytometry | Cat# A21207 |
